# De novo damaging coding mutations are strongly associated with obsessive-compulsive disorder and overlap with autism

**DOI:** 10.1101/127712

**Authors:** Carolina Cappi, Melody E. Oliphant, Zsanett Péter, Gwyneth Zai, Catherine A. W. Sullivan, Abha R. Gupta, Ellen J. Hoffman, Manmeet Virdee, A. Jeremy Willsey, Roseli G. Shavitt, Euripedes C. Miguel, James L. Kennedy, Margaret A. Richter, Thomas V. Fernandez

## Abstract

Obsessive-compulsive disorder (OCD) is a debilitating developmental neuropsychiatric disorder with a genetic risk component, yet identification of high-confidence risk genes has been challenging. We performed whole-exome sequencing in 222 OCD parent-child trios (184 trios after quality control), finding strong evidence that de novo likely gene disrupting and predicted damaging missense variants contribute to OCD risk. Together, these de novo damaging variants are enriched in OCD probands (RR 1.52, p=0.0005). We identified two high-confidence risk genes, each containing two de novo damaging variants in unrelated probands: *CHD8 (Chromodomain Helicase DNA Binding Protein 8)* and *SCUBE1 (Signal Peptide, CUB Domain And EGF Like Domain Containing 1)*. Based on our data, we estimate that 34% of de novo damaging variants seen in OCD contribute to risk, and that de novo damaging variants in approximately 335 genes contribute to risk in 22% of OCD cases. Furthermore, genes harboring de novo damaging variants in OCD are enriched for those reported in neurodevelopmental disorders, particularly autism spectrum disorders. An exploratory network analysis reveals significant functional connectivity and enrichment in canonical pathways related to immune response.

**SIGNIFICANCE STATEMENT:** Decades of genetic studies in obsessive-compulsive disorder (OCD) have yet to provide reproducible, statistically significant findings. Following an approach that has led to tremendous success in gene discovery for several neuropsychiatric disorders, here we report findings from DNA whole-exome sequencing of patients with OCD and their parents. We find strong evidence for the contribution of spontaneous, or de novo, predicted-damaging genetic variants to OCD risk, identify two high-confidence risk genes, and detect significant overlap with genes previously identified in autism. These results change the status quo of OCD genetics by identifying novel OCD risk genes, clarifying the genetic landscape of OCD with respect to de novo variation, and suggesting underlying biological pathways that will improve our understanding of OCD biology.

## INTRODUCTION

Obsessive-compulsive disorder (OCD) is an often-disabling developmental neuropsychiatric disorder with onset typically during adolescence or young adulthood and a lifetime prevalence of 1.5-2.5% (1-5). Obsessions are intrusive thoughts, images, or urges experienced as irrational, excessive, and accompanied by anxiety or discomfort. Compulsions are behaviors undertaken to mitigate obsessions or subjective feelings (i.e., the need to relieve a tactile sensation or achieve a “just right” feeling); they are usually repetitive, stereotyped, and excessive (6, 7). The anxiety or distress associated with obsessions and compulsions and the time spent on them are sources of lifelong morbidity in OCD, having profound negative effects on both patients’ and families’ quality of life. Symptoms can be so disabling that the World Health Organization has ranked OCD among the 10 most debilitating disorders of any kind, in terms of lost earnings and diminished quality of life (8, 9). Furthermore, OCD has been linked to significantly increased mortality, even after controlling for comorbid psychiatric conditions, which can occur in up to 75% of cases (10, 11). Treatment-refractory disease is common, with about 40% of patients resistant to current pharmacological and psychotherapeutic treatments, and untreated OCD generally persists and becomes chronic (12, 13). The causes and underlying pathophysiology of OCD are not well understood, which has limited the development of new treatments and interventions. For these reasons, there is an urgent need for more research to elucidate OCD risk factors and disease mechanisms.

Twin and family studies provide strong evidence for a substantial genetic contribution to OCD risk, with modern estimates of heritability around 40-50% (14-17), yet progress in identifying risk genes has been slow. Decades of linkage, common-variant candidate gene association studies, and more recent genome-wide association studies in OCD (18-20) have yielded few reproducible associations and therefore have provided limited insights into disease biology. Further efforts are clearly needed to identify specific OCD risk variants and to confirm vulnerability pathways by modern genome-wide and comprehensive variant discovery approaches.

In contrast, genetic research into several other neuropsychiatric disorders has seen great progress in recent years, partly attributable to an increasing effort to evaluate the contribution of rare genetic sequence variation, especially de novo variants (arising spontaneously in the parental germ cells or in a zygote shortly after conception). This approach has shown great success for systematic risk gene discovery in other genetically complex neuropsychiatric disorders (21-23), particularly autism spectrum disorders (24-27). While an individual rare variant is unlikely to explain a sizeable fraction of disease risk in the context of a heterogeneous genetic architecture, concurrent investigations of multiple genes implicated by rare sequence and structural variation highlight convergence toward a limited number of important underlying biological mechanisms (28). Therefore, there is a proven avenue for risk gene discovery in complex neuropsychiatric disorders that has yet to be fully leveraged in OCD.

Following these previous studies in other disorders and our pilot study suggesting a role for de novo single nucleotide variants (SNVs) in OCD risk (29), we performed whole-exome sequencing (WES) in 222 OCD parent-child trios to identify de novo SNVs and insertion-deletion variants (indels). In 184 OCD trios passing quality control, we find strong evidence for the contribution of de novo likely gene disrupting (LGD; disruption of a stop codon, canonical splice site, or a frameshift indel) as well as predicted damaging missense (Mis-D) variants to OCD. Furthermore, we identified two high-confidence candidate risk genes based on observing gene-level recurrence of de novo damaging (LGD + Mis-D) variants in unrelated probands: *CHD8 (Chromodomain Helicase DNA Binding Protein 8)* and *SCUBE1 (Signal Peptide, CUB Domain And EGF Like Domain Containing 1)*. We estimated that 22% of OCD cases will harbor a de novo damaging SNV or indel mediating OCD risk, and that there are approximately 335 genes affected by such variants contributing to the risk. Finally, we detected significant overlap between genes with damaging de novo variants in OCD and autism.

## RESULTS

### Damaging de novo SNVs and indels are associated with OCD risk

Exome sequencing was performed on 222 OCD parent-child trios. WES data from 855 unaffected trios, already sequenced from the Simons Simplex Collection, were pooled with our OCD trios for joint variant calling. After quality control methods, our sample size for a burden analysis was 184 OCD and 777 unaffected trios (Figure 1, Table 1, Table S1). To compare the de novo mutation rates between cases and controls, we limited our analysis to loci with at least 20x coverage in all members of a trio, as this was our pre-defined threshold for calling a de novo variant (see Methods). Based on our OCD pilot study (29) and work in other neurodevelopmental disorders (22, 24, 26, 30), we expected to find an enrichment of de novo LGD variants (stop codon, frameshift, or canonical splice-site variants) in OCD probands versus controls. We found a statistically significant increased rate of de novo LGD variants in OCD cases, confirming our hypothesis (rate ratio [RR] 1.93, 95% Confidence Interval [CI] 1.19-3.09, p=0.01). Furthermore, de novo missense variants predicted to be damaging by PolyPhen2 (Mis-D; Polyphen2 HDIV score ≥0.957) were also over-represented in OCD probands (RR 1.43, CI 1.13-1.80, p=0.006). Taken together, damaging de novo coding variants (LGD and Mis-D) occur more often in OCD probands versus controls (RR 1.52, CI 1.23-1.86, p=0.0005). We did not detect a difference in mutation rates for de novo synonymous variants (RR 0.99, CI 0.75-1.31, p=0.5). (See Table 1, Figure 2A)

**Figure 1.**
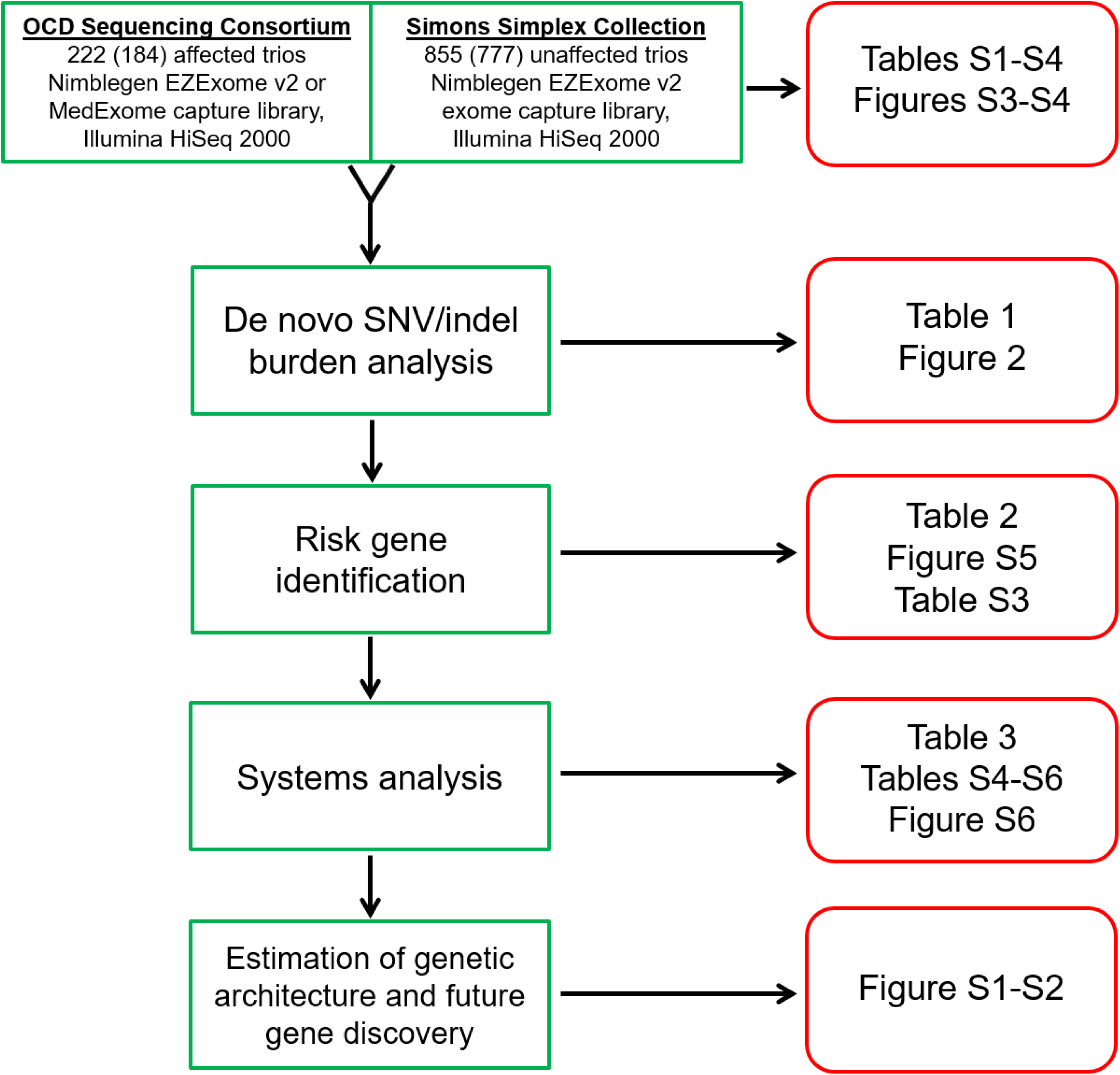
Study summary. We performed whole exome sequencing on 222 OCD and 855 control parent-child trios. After quality control, 184 OCD and 777 control trios remained for subsequent analyses. A burden analysis compared the rates of de novo single nucleotide (SNVs) and insertion-deletion (indel) variants between cases and controls. Next, we used the TADA-Denovo algorithm to assess the significance of gene-level recurrence of de novo damaging variants in our OCD group, identifying two high-confidence risk genes. Exploratory network, pathway, and cross-disorder analyses were then performed using genes harboring de novo damaging variants in our OCD subjects. Finally, based on the number of de novo damaging variants in OCD versus controls, we estimated the number of genes contributing to OCD risk, and used this estimate to predict future risk gene discovery as additional OCD parent-child trios are studied by exome sequencing.

**Figure 2.**
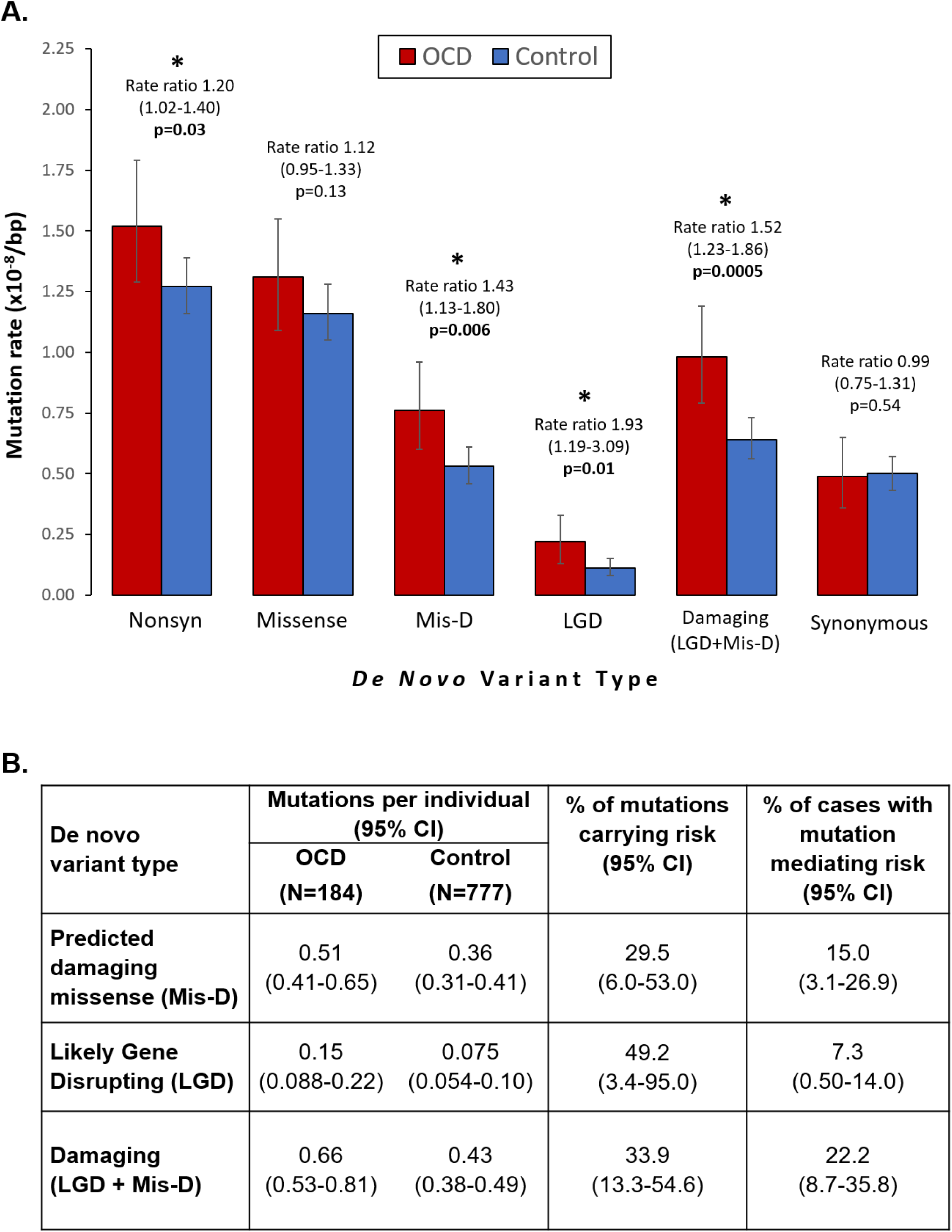
De novo damaging variants are associated with OCD risk. **(A)** Bar chart comparing the rates of de novo mutation types between OCD cases (red) and controls (blue). Comparisons are between per base pair (bp) mutation rates, considering only those “callable” loci in each family and cohort that meet required sequencing depth and quality scores to support high confidence de novo variant calling. Mutation rates were compared using a one-tailed rate ratio test. Statistically significant comparisons (p<0.05) are marked with asterisks. Error bars show 95% confidence intervals. **(B)** For the enriched classes of de novo variants, we quantified their contribution to OCD risk in two ways. First, we estimated the percentage of observed variants carrying risk by dividing the difference in rates (estimated coding variants per individual, see Table 1 and Methods) by the rate in OCD. Second, we estimated the percentage of cases with a mutation mediating risk by subtracting the proportion of controls carrying a mutation from the proportion in OCD probands carrying a mutation.

**Table 1.** Distribution of de novo variants in OCD cases and controls.

| De novo variant type<sup>a</sup> | Variant counts |  | Mutation rate (x10<sup>-8</sup>) per bp (95% CI)<sup>j</sup> |  | Estimated coding variants per individual (95% CI)<sup>k</sup> |  | Rate ratio (95% CI) | p-value<sup>l</sup> |
| --- | --- | --- | --- | --- | --- | --- | --- | --- |
|  | OCD (N=184) | Control (N=777) | OCD (N=184) | Control (N=777) | OCD (N=184) | Control (N=777) |  |  |
| All<sup>b</sup> | 207 | 701 | 2.02<br>(1.75-2.31) | 1.80<br>(1.67-1.94) | 1.37<br>(1.18-1.56) | 1.22<br>(1.13-1.31) | 1.12<br>(0.95-1.31) | 0.11 |
| Coding<sup>c</sup> | 200 | 662 | 2.06<br>(1.78-2.36) | 1.80<br>(1.67-1.95) | 1.39<br>(1.20-1.60) | 1.22<br>(1.13-1.32) | 1.14<br>(0.99-1.30) | 0.06 |
| Synonymous SNV | 48 | 182 | 0.49<br>(0.36-0.65) | 0.50<br>(0.43-0.57) | 0.33<br>(0.24-0.44) | 0.34<br>(0.29-0.39) | 0.99<br>(0.75-1.31) | 0.54 |
| Nonsynonymous<sup>d</sup> | 148 | 467 | 1.52<br>(1.29-1.79) | 1.27<br>(1.16-1.39) | 1.03<br>(0.87-1.21) | 0.86<br>(0.78-0.94) | **<u>1.20</u>**<br>**<u>(1.02-1.40)</u>** | **<u>0.03</u>** |
| All Missense (Mis) | 127 | 426 | 1.31<br>(1.09-1.55) | 1.16<br>(1.05-1.28) | 0.89<br>(0.74-0.20) | 0.78<br>(0.71-0.87) | 1.12<br>(0.95-1.33) | 0.13 |
| Mis-D<sup>e</sup> | 74 | 195 | 0.76<br>(0.60-0.96) | 0.53<br>(0.46-0.61) | 0.51<br>(0.41-0.65) | 0.36<br>(0.31-0.41) | **<u>1.43</u>**<br>**<u>(1.13-1.80)</u>** | **<u>0.006</u>** |
| Mis-P<sup>f</sup> | 18 | 79 | 0.19<br>(0.11-0.29) | 0.22<br>(0.17-0.27) | 0.13<br>(0.074-0.20) | 0.15<br>(0.12-0.18) | 0.86<br>(0.53-1.34) | 0.76 |
| Mis-B<sup>g</sup> | 33 | 147 | 0.34<br>(0.23-0.48) | 0.40<br>(0.34-0.47) | 0.23<br>(0.16-0.32) | 0.27<br>(0.23-0.32) | 0.85<br>(0.60-1.17) | 0.83 |
| Likely Gene Disrupting (LGD)<sup>h</sup> | 21 | 41 | 0.22<br>(0.13-0.33) | 0.11<br>(0.080-0.15) | 0.15<br>(0.088-0.22) | 0.074<br>(0.054-0.10) | **<u>1.93</u>**<br>**<u>(1.19-3.09)</u>** | **<u>0.01</u>** |
| Damaging (LGD + Mis-D) | 95 | 236 | 0.98<br>(0.79-1.19) | 0.64<br>(0.56-0.73) | 0.66<br>(0.53-0.81) | 0.43<br>(0.38-0.49) | **<u>1.52</u>**<br>**<u>(1.23-1.86)</u>** | **<u>0.0005</u>** |
| LGD SNV | 14 | 20 | 0.14<br>(0.079-0.24) | 0.055<br>(0.033-0.084) | 0.095<br>(0.053-0.16) | 0.037<br>(0.022-0.057) | **<u>2.64</u>**<br>**<u>(1.39-4.93)</u>** | **<u>0.006</u>** |
| LGD frameshift indel | 7 | 21 | 0.072<br>(0.029-0.15) | 0.057<br>(0.035-0.088) | 0.049<br>(0.020-0.10) | 0.039<br>(0.024-0.060) | 1.28<br>(0.53-2.72) | 0.37 |
| **Nonframeshift indel** | 2 | 5 | 0.021<br>(0.0025-0.074) | 0.014<br>(0.0044-0.032) | 0.014<br>(0.00017-0.050) | 0.0095<br>(0.0030-0.022) | 1.51<br>(0.21-7.28) | 0.44 |
| **Unknown<sup>i</sup>** | 2 | 8 | 0.021<br>(0.0025-0.074) | 0.022<br>(0.0094-0.043) | 0.014<br>(0.00017-0.050) | 0.015<br>(0.0064-0.029) | 0.94<br>(0.14-3.88) | 0.65 |

### Damaging de novo SNVs and indels contribute to OCD risk in 22% of cases

Next, we estimated the fraction of observed de novo mutations that contribute to OCD risk, based on our dataset. By dividing the de novo mutation rate difference between cases and controls by the rate in cases, we estimate that 49.2% (CI 3.4-95.0%) of de novo LGD and 29.5% (CI 6.0-53.0%) of de novo Mis-D mutations contribute to OCD risk. As a group, we estimate that 33.9% (CI 13.3-54.6%) of damaging (LGD + Mis-D) de novo mutations contribute to OCD risk (Figure 2B).

We also used our data to estimate the proportion of cases harboring a de novo mutation contributing to OCD risk. By subtracting the percentage of controls from the percentage of OCD probands with at least one de novo mutation, we estimate that 15.0% (CI 3.1-26.9%) have a de novo Mis-D mutation and 7.3% (CI 0.50-14.0%) have a de novo LGD mutation mediating OCD risk. As a group, we estimate that 22.2% (CI 8.7-35.8%) of cases have a damaging de novo mutation contributing to OCD risk (Figure 2B).

### Recurrent damaging de novo variants identify two candidate risk genes

Having established that de novo damaging variants occur more frequently in OCD probands, we next asked whether these variants cluster within specific genes. We identified three genes with multiple (≥2) de novo LGD or Mis-D variants in unrelated probands. Using TADA-Denovo (31) and previously established false discovery rate (FDR) thresholds, two of these genes met criteria for high-confidence risk genes (q<0.1): *SCUBE1* (*Signal Peptide, CUB Domain And EGF Like Domain Containing 1*; q=0.040) and *CHD8* (*Chromodomain Helicase DNA Binding Protein 8*; q=0.043) (Table 2). A third gene, *TTN* (*Titin*), did not meet this threshold (q=0.47) (Table 2).

**Table 2.** Risk gene discovery in OCD.

| Gene | # LGD | # Mis-D | p-value | q-value (FDR) |
| --- | --- | --- | --- | --- |
| *SCUBE1* | 1 | 1 | $2.1 \times 10^{-6}$ | 0.040 |
| *CHD8* | 1 | 1 | $2.5 \times 10^{-6}$ | 0.043 |
| *TTN* | 0 | 3 | 0.0005 | 0.47 |

### Approximately 335 genes contribute to OCD risk

Based on OCD proband vulnerability to de novo damaging variants in our dataset, we used two methods to estimate the number of genes contributing to OCD risk. Using a maximum likelihood estimation (MLE) method (32), we determined the most likely number of genes to be 335 (Figure S1). This agrees with an alternate method based on the “unseen species problem” (33); the estimated number of OCD risk genes using this alternate method is 317 (95% CI 190-454).

Next, we used the estimated number of OCD risk genes (n=335) to predict the likely future gene discovery yield as additional OCD trios are investigated by WES. Based upon 10,000 simulations at each cohort size, we predict discovery of the following numbers of risk genes as we sequence more OCD parent-child trios: 24 probable risk genes, including 11 high-confidence risk genes (24 / 11 genes) at 500 trios; 77 / 40 genes at 1,000 trios; 202 / 113 genes at 2,000 trios; 323 / 189 genes at 3,000 trios (Figure S2).

### Overlap with ASD and CHD8 target genes

Using DNENRICH (34), we found significant overlap between genes harboring de novo damaging variants in OCD (n=89, excludes occurrences in controls) and several gene sets from the literature (Table 3, Table S4). Our OCD genes were significantly enriched for genes harboring de novo nonsynonymous (LGD, missense) variants in autism (ASD), genes achieving TADA q<0.1 in ASD, genes with genome-wide significant statistical evidence for association with developmental disorders, and genes that are targets of CHD8 in the developing human brain. There was no significant enrichment for genes harboring de novo variants in intellectual disability (ID) or schizophrenia (SCZ), no enrichment for de novo synonymous variants in any disorder, and no enrichment for any class of de novo variation in unaffected siblings in the SSC (Table 3, Table S4).

**Table 3.** Overlap between OCD de novo damaging mutations and gene sets.

| Comparison<br>gene set<sup>a</sup> | LGD |  |  |  | Missense |  |  |  | Synonymous |  |  |  |
| --- | --- | --- | --- | --- | --- | --- | --- | --- | --- | --- | --- | --- |
|  | Obs<sup>b</sup> | Exp<sup>c</sup> | O/E<sup>d</sup> | P<sup>e</sup> | Obs | Exp | O/E | P | Obs | Exp | O/E | P |
| ASD | 9 | 4.55 | 1.98 | **0.038** | 25 | 17.94 | 1.39 | **0.043** | 9 | 8.06 | 1.12 | 0.41 |
| SCZ | 1 | 1.00 | 1.00 | 0.64 | 7 | 5.40 | 1.30 | 0.29 | 0 | 1.84 | 0 | 1 |
| DD | 2 | 2.77 | 0.72 | 0.77 | 5 | 6.50 | 0.77 | 0.79 | 4 | 2.27 | 1.76 | 0.19 |
| TD | 0 | 0.26 | 0 | 1 | 2 | 1.90 | 1.05 | 0.57 | 2 | 0.77 | 2.61 | 0.18 |
| ID | 0 | 0.49 | 0 | 1 | 1 | 1.28 | 0.78 | 0.72 | 0 | 0.43 | 0 | 1 |
| Unaffected | 2 | 1.35 | 1.48 | 0.39 | 5 | 8.98 | 0.56 | 0.95 | 2 | 4.20 | 0.48 | 0.93 |
|  | **Obs** | **Exp** | **O/E** | **P** |  |  |  |  |  |  |  |  |
| DD significant<sup>f</sup> | 4 | 0.84 | 4.76 | **0.010** |  |  |  |  |  |  |  |  |
| ASD – q<0.1<sup>g</sup> | 3 | 0.65 | 4.62 | **0.028** |  |  |  |  |  |  |  |  |
| CHD8 brain<sup>h</sup> | 20 | 13.02 | 1.54 | **0.030** |  |  |  |  |  |  |  |  |

<sup>a</sup>Comparisons for autism (ASD), schizophrenia (SCZ), developmental disorders (DD), Tourette’s disorder (TD), intellectual disability (ID), and unaffected siblings are between genes harboring de novo LGD, missense, or synonymous variants in these phenotypes and those harboring damaging (LGD or Mis-D) variants in OCD (n=89). Gene lists and their references are in Table S4. <sup>b</sup>Observed number of genes overlapping between sets. <sup>c</sup>Expected number of genes overlapping between sets, determined by 100,000 random mutation set simulations using DNENRICH (Fromer et al., 2014). <sup>d</sup>O/E, observed divided by expected number of genes overlapping between sets. <sup>e</sup>P-value is one-sided under a binomial model of greater than expected hits per gene set, calculated by DNENRICH using 100,000 permutations. <sup>f</sup>93 genes with genome-wide significant statistical evidence for association with developmental disorders (Deciphering Developmental Disorders Study, 2017). <sup>g</sup>65 ASD genes with False Discovery Rate (q) < 0.1 by TADA, considering data across exome sequencing studies (Sanders et al., 2015). <sup>h</sup>Genes in human brain with promoters targeted by CHD8 (Cotney et al., 2015). See also Tables S2 and S4.

### Exploratory pathway and network analyses

Using our list of genes harboring de novo damaging variants in OCD (n=89), we performed exploratory analyses to determine shared underlying canonical pathways and functional connectivity. Using the GeNets algorithm, OCD genes mapping onto a meta-network displayed significantly more connectivity than expected by chance (p=0.026), and 68 additional “tier 1” candidate genes were predicted, based on their high connectivity to our input genes (Figure S6, Table S5). Our input gene list is significantly enriched for canonical pathways related to immune response, particularly the complement system, which was a top result from two pathway analysis tools, MetaCore and IPA. Other significant canonical pathways include granulocyte-macrophage colony stimulating factor (GM-CSF) signaling, neurotrophin/tyrosine kinase signaling, B cell receptor signaling, and focal adhesion kinase signaling (Table S6).

## DISCUSSION

By whole-exome sequencing of OCD parent-child trios, we have demonstrated a strong association between de novo damaging (LGD and Mis-D) coding variants and OCD cases (Table 1, Figure 2). As seen in studies of other neurodevelopmental disorders, these results can be leveraged to systematically identify OCD risk genes. In the current study, two genes, *CHD8* and *SCUBE1*, have a FDR q<0.1, meeting criteria for high-confidence association with OCD (Table 2).

*SCUBE1* has not been extensively studied. While it is expressed in the developing brain and nervous system (35, 36), functional studies to date have focused mostly on its potential role in platelet activation and adhesion (37, 38). A study in mice has shown downregulated *SCUBE1* expression in response to inflammatory stimuli (36), but this gene has not yet been implicated in disorders of the brain or nervous system. Interestingly, increased levels of pro-inflammatory markers have been reported in several studies of children and adolescents with OCD (39-42).

On the other hand, there are several recent and ongoing studies of *CHD8*, a gene that has emerged as having the strongest association with autism spectrum disorder via the identification of multiple de novo LGD variants in unrelated parent-child trios (Figure S5) (33, 43-45). *CHD8* is highly expressed in the developing brain (46). It encodes an ATP-dependent chromatin remodeler that binds to tri-methylated histone H3 lysine 4, a post-translational histone modification present at active promoters (47-49). Loss of *CHD8* function appears to contribute to autism pathology by disrupting the expression of its target genes, which are themselves enriched for high confidence autism risk genes (46). While OCD subjects with de novo damaging *CHD8* variants in our study do not meet any diagnostic criteria for autism, this finding suggests there may be overlapping biological mechanisms between the two disorders, and leads us to hypothesize that genes regulated by *CHD8* may similarly be enriched for OCD risk genes. Indeed, we see significant overlap between our OCD genes and ASD genes, as well as *CHD8* gene targets mapped in the developing human brain (46) (Table 3).

Based on data from this study, we estimate that 34% of de novo damaging mutations seen in OCD carry risk (Figure 2B) and that 335 genes confer risk in 22% of patients (Figure 2B). Given our OCD sample size, the 95% confidence intervals around these contribution estimates are wide, and need refinement by continued sequencing of OCD trios.

Mindful of the fact that more than half of genes harboring de novo damaging variants in our study may not be true risk genes, we consider our pathway and network analyses as exploratory at this stage. Nevertheless, we see preliminary evidence that genes identified by de novo damaging variants in OCD are functionally connected to a greater degree than expected by chance (Figure S6, Table S5). Furthermore, these genes may be enriched in immunological and cell signaling canonical pathways (Table S6), consistent with our pilot study of exome sequencing in 20 OCD trios (29). These analyses should be repeated as more high-confidence OCD risk genes are identified.

While not rising to the level of a high-confidence risk gene in this study, it is notable that we identified an OCD de novo damaging (Mis-D) variant in *DLGAP1* (*discs, large homolog- associated protein 1*). In a genome-wide association study by the International OCD Foundation Genetic Collaborative (IOCDF-GC), the lowest p-values for their case-control analysis were found for two SNPs located within *DLGAP1* (2.49×10^−6^, 3.44x10^−6^) (20). A subsequent GWAS by the OCD Collaborative Genetics Study (OCGAS) identified a SNP nearby this gene with a prominent signal (p=2.67×10^−4^) (18). Furthermore, a rare paternally-inherited duplication in *DLGAP1* was recently reported in a child with OCD, Tourette syndrome, and anxiety (50). *DLGAP1* is a member of the neuronal postsynaptic density complex and is in the same family as *DLGAP3* (*SAPAP3*), a gene associated with OCD-like behaviors in a knockout mouse model (51). Therefore, evidence is beginning to converge on this gene as one of great interest in OCD genetics.

Successful gene discovery by leveraging gene-level recurrent de novo variation in autism, where over 65 genes have now been identified (24, 26, 27), and the results presented here for OCD, strongly reinforce the value of continuing WES in larger cohorts of OCD parent-child trios. Our models predict that by increasing the sample size of this study to 500 trios, we will gain 9 additional high-confidence risk genes and 22 probable risk genes. Further increasing to 1,000 trios will yield a total of 40 high-confidence and 77 probable risk genes. Discovering risk genes will change the status quo in OCD genetics by allowing new studies in model systems (e.g. animal models, induced pluripotent stem cells) and network analyses. Such studies will provide insights into OCD pathophysiology that are critical prerequisites for the discovery of novel therapeutic targets to alleviate the suffering of those with OCD.

## MATERIALS AND METHODS

### Subjects

This study was approved by the local institutional review boards of all participating institutions, and appropriate informed consent was obtained from participating subjects. 222 parent-child trios (139 male, 83 female), consisting of offspring meeting criteria for the diagnosis of obsessive-compulsive disorder, as defined by the Diagnostic and Statistical Manual for Mental Disorders (DSM-IV-TR or DSM-5) (52, 53), and their unaffected parents, were recruited for DNA sequencing. Trios were recruited at three sites: the University of São Paulo School of Medicine Obsessive-Compulsive Spectrum Disorders Program (42 trios), Centre for Addiction and Mental Health and the Frederick W. Thompson Anxiety Disorders Centre at the Sunnybrook Health Sciences Centre in Toronto (77 trios), and Yale University School of Medicine (61 trios). Additionally, we included 42 trios with OCD and chronic tics that were recruited for a separate study by TIC Genetics (54, 55). Other diagnostic criteria included: onset of symptoms prior to age 18 years; no previously diagnosed neurological disorder, intellectual disability, psychotic disorder, or OCD occurring exclusively in the context of depression; no known history of OCD in first degree relatives. Final diagnostic status was assigned based on the consensus of an experienced interviewer and a psychiatrist or psychologist after independent review and administration of a structured diagnostic interview. We prioritized the study of simplex OCD trios to increase the likelihood of detecting de novo sequence and structural variants. Available phenotype information, including gender and parental age, is included in Table S1.

### Whole-exome sequencing (WES)

Exome capture and sequencing of blood-derived DNA from 222 affected children and their parents (666 samples total) were performed at the Yale Center for Genomic Analysis (YCGA), using the NimbleGen SeqCap EZExomeV2 (109 trios) or MedExome (113 trios) capture libraries (Roche NimbleGen, Madison, WI, USA) and the Illumina HiSeq 2000 platform (74 bp paired-end reads; Illumina, San Diego, CA). We multiplexed six samples during each capture reaction and sequencing lane, pooling parents and probands when possible. WES data from 855 unaffected parent-child trios (2565 samples total) were obtained from the Simons Simplex Collection via the NIH Data Archive (https://ndar.nih.gov/edit_collection.html?id=2042). These children and their parents have no evidence of autism spectrum or other neurodevelopmental disorders (56). Like our OCD samples, control WES was from blood-derived DNA and sequenced on the Illumina HiSeq 2000 sequencing platform after capture with the NimbleGen SeqCap EZExomeV2 library.

### Sequence alignment and variant calling

Alignment and variant calling of the sequencing reads followed the latest Genome Analysis Toolkit (GATK) (57) Best Practices guidelines. Reads were aligned using BWA-mem (58) to the b37 human reference sequence with decoy sequences. Picard’s MarkDuplicates tool was used to mark PCR duplicates (https://broadinstitute.github.io/picard/). A target bed file was created by taking the intersection of the EZExomeV2 and MedExome target regions. GATK was used to realign indels, recalibrate quality scores, and generate GVCF files for each sample using the HaplotypeCaller tool. All samples were called jointly using GATK’s GenotypeGVCFs tool, variant score recalibration was applied to the called variants, and all variant call data was written to a VCF file. This pipeline uses GATK’s Best Practices parameters and the default parameters for BWA and Picard. Only passing variants were used in downstream analyses. Variants were annotated against the RefSeq hg19 gene definitions and multiple external databases of variant population frequency, conservation scores, variation intolerance, mutation severity, and predicted functional effects using ANNOVAR (59).

### Quality control and de novo variant calling

Relatedness statistics were calculated based on the method of Manichaikul et al. (60), implemented in VCFtools v0.1.14.10 (61). Trios were omitted if expected family relationships were not confirmed or if there were unexpected relationships within or between families. Trios were omitted if > 5 de novo variants were observed. PLINK/SEQ (34) (i-stats; https://psychgen.u.hpc.mssm.edu/plinkseq/stats.shtml), PicardTools, and GATK DepthOfCoverage tools were used to generate quality metrics (Table S1). To identify outliers that might confound our case-control analysis, we performed principal components analysis (PCA) using this data. A scree plot determined the number of principal components accounting for the greatest proportion of variance, and we removed trios with family members falling more than three standard deviations from the mean in any of these principal components. See Figure S3, Table S1, and Supplementary Methods.

We used stringent thresholds for identifying de novo mutations because DNA from control subjects (recruited by the Simons Foundation Autism Research Initiative) was not available for confirmation by Sanger sequencing. De novo variants were called using an in-house script that required the following: child is heterozygous for a variant with alternate allele frequency between 0.3 and 0.7 in the child and < 0.05 in the parents, sequencing depth (DP) ≥ 20 in all family members at the variant position, alternate allele depth (AD) ≥ 5, observed allele frequency (AC) < 0.01 (1%) among all cases and controls, mapping quality (MQ) ≥ 30. False positive calls were removed by in silico visualization. We performed Sanger sequencing on the probands and parents for 383 putative de novo variants from this and another exome sequencing project using the same calling thresholds; 370 were confirmed, resulting in 96.6% specificity.

### Mutation rate analysis

Within each cohort, we calculated the rate of de novo mutations per base pair. For accurate rate calculation, we first determined the number of “callable” base pairs per family using the GATK DepthOfCoverage tool. We considered only bases covered at ≥ 20x in all family members, with base quality ≥ 20, and map quality ≥ 30; these thresholds match those required for GATK and de novo variant calling. For each cohort, we summed the “callable” base pairs in every family and used this number as the denominator for de novo rate calculations. The resulting rate was divided by two to give haploid rates. Confidence intervals were calculated using the *pois.conf.int (pois.exact)* function from the epitools v0.5-9 package in R. We compared de novo mutation rates in cases versus controls (burden analysis) using a one-tailed rate ratio test in R (https://cran.r-project.org/package=rateratio.test), considering only those variants present with a frequency of <0.01 in the ExAC v0.3.1 database (62). See Table 1, Table S1, and Supplementary Methods.

### TADA analysis

Prior exome analyses demonstrated that the observation of even a small number of rare de novo mutations in the same gene among unrelated individuals can provide considerable statistical power to establish association (33). We used the Transmitted And De novo Association (TADA-Denovo) test as a statistical method for risk gene discovery based on gene-level recurrence of de novo mutations within the classes of variants that we found enriched in OCD (27, 31). Parameter calculations and a detailed description of the method are given in Supplementary Methods. In summary, TADA-Denovo generates random mutational data based on each gene’s specified mutation rate to determine null distributions, then calculates a p-value and a false discovery rate (FDR) q-value for each gene using a Bayesian “direct posterior approach.” A low q-value represents strong evidence for OCD association. See Table 2, Table S3, and Supplementary Methods.

### Estimation of number of risk gene

We first used a maximum likelihood estimation (MLE) method to estimate the number of genes contributing risk to OCD, based on vulnerability to de novo damaging variants (32). For every number of risk genes from 1 to 2,500, we simulated 95 variants (the number of damaging de novo variants observed in probands in our case-control burden analysis). Variant simulations were performed 100,000 times at each number of risk genes. Following each simulation, a percentage of variants was randomly assigned to the risk genes. The percentage of variants assigned to risk genes was determined by the fraction of de novo damaging variants estimated to carry OCD risk, and variant simulations were weighted by gene size and GC content (31). We then counted the number of risk and non-risk genes containing two variants and the number containing three or more variants. The frequency of concordance between our simulated and observed data was calculated. A curve was plotted to show the concordance frequency (y-axis) at each assumed number of risk genes (x-axis), and the peak was taken as the estimate of the most likely number of risk genes (32). See Figure S1 and Supplementary Methods.

Next, we used an alternate method for estimating the number of risk genes, using a statistical method based on the “unseen species” problem (33). This method uses the frequency and number of observed variant types (or species) to infer how many species are present in the population. See Supplementary Methods for details of these calculations.

### Estimation of future risk gene discovery

We used the estimated number of OCD risk genes to predict the likely future gene discovery yield as additional OCD trios are investigated by WES. Fixing the gene number at 335 (from MLE estimate above), we varied the cohort size (from 25 to 3000, in increments of 25). At each cohort size, we simulated a number of variants matching the observed mutation rate in OCD probands. Simulated variants were randomly assigned to the risk genes. The percentage of variants assigned to risk genes was determined by the fraction of de novo damaging variants estimated to carry OCD risk, and variant simulations were weighted by gene size and GC content (31). At each cohort size, 10,000 simulations were performed. LGD and Mis-D variants were generated separately. Simulated variants were then combined and given as input to the TADA-Denovo algorithm, using the same parameters described above for the observed data. The number of high confidence (q<0.1) and probable (q<0.3) risk genes were recorded and plotted using polynomial regression fitting; this regression model allows prediction of the number of genes identified at a specified cohort size. See Figure S2 and Supplementary Methods.

### Gene set overlap

We used DNENRICH (34) (https://psychgen.u.hpc.mssm.edu/dnenrich/) to test whether OCD genes harboring de novo damaging mutations (89 genes; excluding two genes, *TTN* and *CACNA1E*, found to harbor de novo damaging variants in control subjects) were significantly enriched among genes identified in autism (ASD), schizophrenia (SCZ), developmental disorders (DD), Tourette’s disorder (TD), and intellectual disability (ID). Gene lists were obtained from a recent cross-disorder study (63) that included de novo single nucleotide and indel variants from multiple exome sequencing studies in ASD (24, 26, 43, 64, 65), SCZ (34, 66-69), and ID (23, 70-72). Three of these studies also included de novo variants present in unaffected siblings (26, 64, 67). DD (21) and TD (54) genes were obtained from recently published WES studies. DNENRICH simulates random mutations while accounting for gene size, trinucleotide context, and mutational effect. We performed 100,000 permutations, comparing the observed and expected overlap with each gene set. Empirical p-values were generated, based on a one-sided enrichment analysis under a binomial model of greater than expected hits per gene set. We tested for overlap between our OCD genes and those in ASD, SCZ, DD, TD, ID, and unaffected siblings that harbored de novo (LGD, nonsynonymous, synonymous) mutations. We also tested for overlap with ASD genes achieving q<0.01 in a recent meta-analysis (27) and DD genes achieving genome-wide statistical significance (21). Finally, given that we identified *CHD8* as an OCD risk gene in our study, we tested for overlap with lists of genes that are targets of CHD8 in human brain (46), See Table 3, Table S4, and Supplementary Methods.

### Exploratory pathway and network analyses

To determine whether all genes harboring de novo damaging variants in OCD are enriched for specific biological pathways, we used the same gene list from our gene set overlap analysis (n=89) to identify the most significant canonical pathways suggested by MetaCore (Thomson Reuters, version 6.30, build 68780, https://portal.genego.com/) and Ingenuity Pathway Analysis (IPA, build version 430520M, content version 31813282, release date 2016-12-05; Ingenuity Systems, http://www.ingenuity.com/). The following default settings were used for MetaCore: Analyze Single Experiment Tool, species Homo sapiens, threshold 0, p-value 1, signals both. The following default settings were used for IPA: Reference set: Ingenuity Knowledge Base (Genes Only); direct and indirect relationships; does not include endogenous chemicals; consider only relationships where species = human and confidence = experimentally observed. See Table S6.

Using the GeNets algorithm (https://apps.broadinstitute.org/genets), we mapped all 89 genes harboring de novo damaging mutations in OCD onto the GeNets Metanetwork v1.0 to determine whether they are functionally connected. The GeNets Metanetwork contains integrated protein-protein interactions from InWeb3 (73), ConcensusPathDB (http://consensuspathdb.org) (74), and 5,057 drug-target interactions (75); the total network size is 530,532 interactions. The GeNets algorithm determines the density of the mapped network (density = number of edges / number of possible edges) and compares this to computed densities for randomly sampled gene sets. An empirically determined p-value is generated. The network is determined to be significantly more connected than random if the density is greater than 95% of the randomly sampled gene sets. Additionally, in the process of mapping our genes onto the Metanetwork, additional candidate genes are predicted, based on their connectivity to our input genes. An overall network connectivity p-value is generated, both with and without these additional predicted candidates. Also, as part of the GeNets analysis, gene “communities” were determined, defined as genes that are more connected to one another than they are to other groups of genes. See Figure S6 and Table S5. All GeNets results for this analysis are also available in interactive form here: https://www.broadinstitute.org/genets#/visualize/58d9425ea4e00291af652379

## ACKNOWLEDGEMENTS

We wish to thank the families who have participated in and contributed to this study. Control subject data were obtained from the NIH-supported National Database for Autism Research (NDAR). NDAR is a collaborative informatics system created by the National Institutes of Health to provide a national resource to support and accelerate research in autism. Dataset identifier: 2042. This manuscript reflects the views of the authors and may not reflect the opinions or views of the NIH or of the Submitters submitting original data to NDAR. Additionally, we wish to thank the Tourette International Collaborative Genetics Study for contributing published genetic data. This study was supported by grants from the Allison Family Foundation [TVF]; FAPESP (process number: 2014/01585-5) and CNPq (process number: 460928/2014-7) [CC]; the Ontario Mental Health Foundation (OMHF) and private donations from the Frederick W. Thompson family [MAR, JLK]. JLK is a Scientific Advisory Board member of AssureRx. JLK has received speaker honoraria and expenses from Eli Lilly and Novartis, consultant honoraria and expenses from Roche, and expenses from AssureRx. MAR has received research support through grants from Roche and speaker honoraria from Lundbeck. TVF has received research support from Shire, the Simons Foundation, and the National Institute of Mental Health; neither funded this project. The authors declare no potential conflict of interest.

## SUPPLEMENTARY METHODS

### Principal Component Analysis (PCA)

PCA was performed on all sequencing quality metrics (Table S1) in R using the following code:

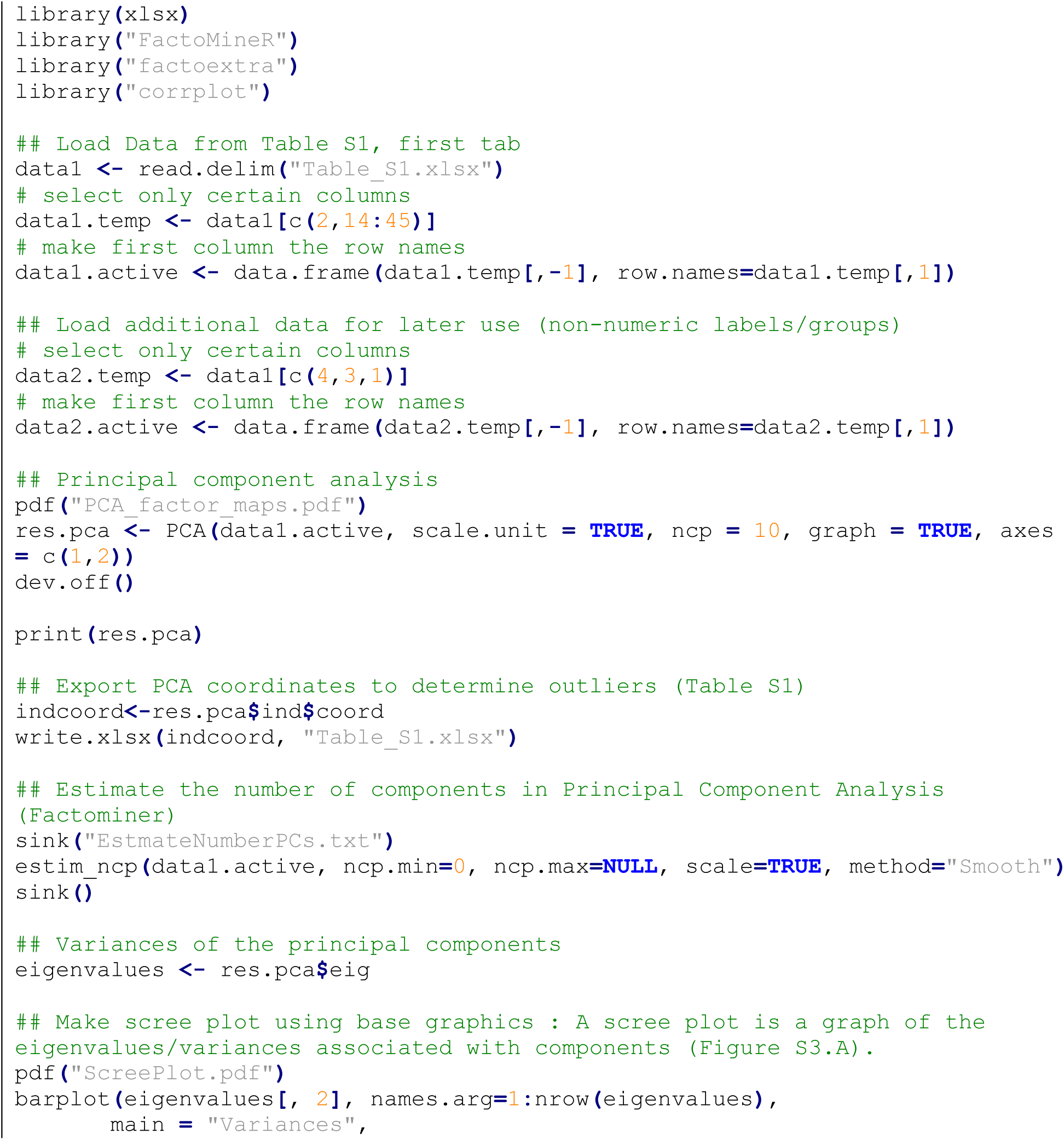

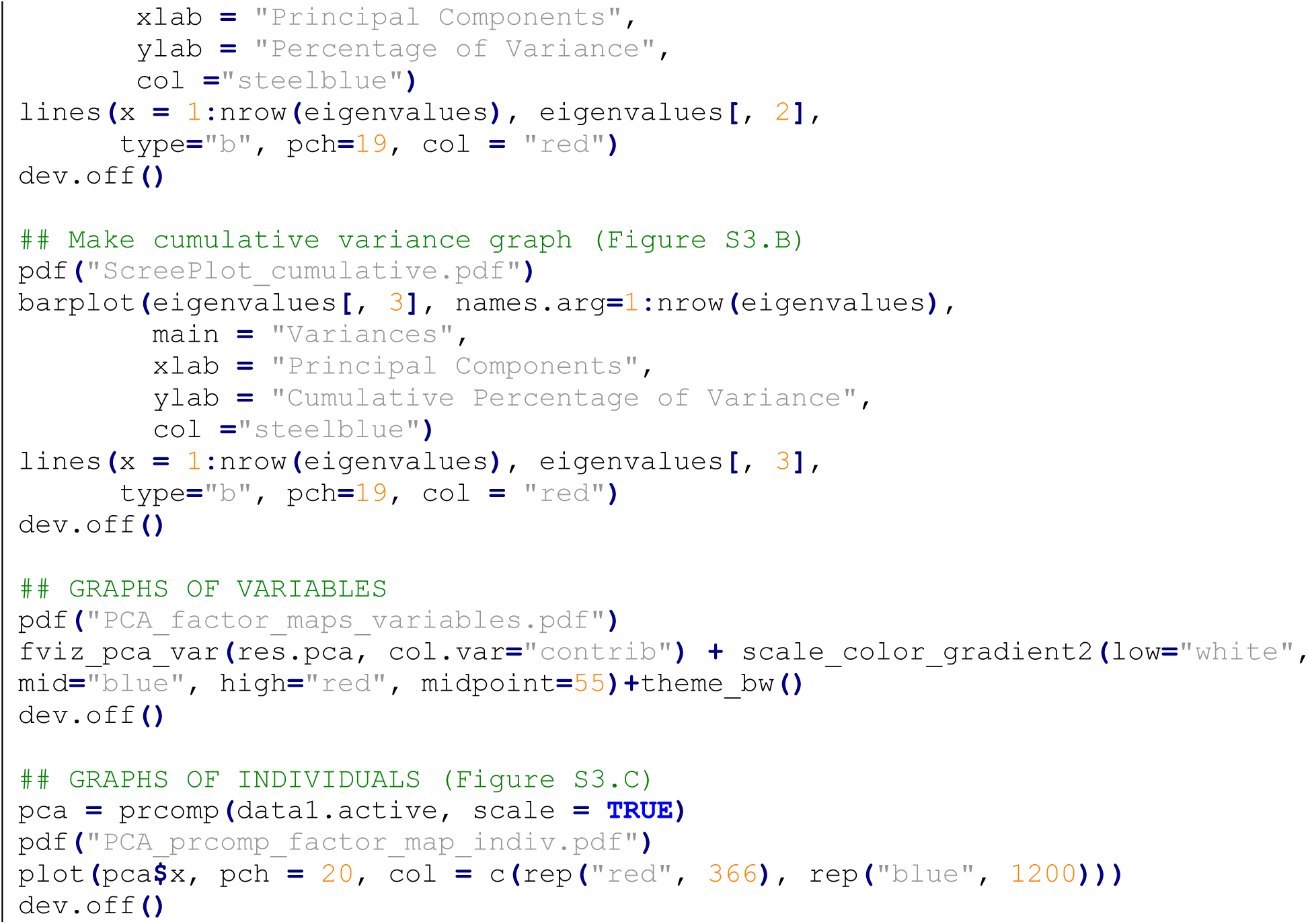

### Calculating “callable” base pairs

Within each cohort, we calculated the rate of de novo mutations per base pair. For accurate rate calculation, we first determined the number of “callable” base pairs per family using the GATK DepthOfCoverage tool. We considered only bases covered at ≥ 20x in all family members, with base quality ≥ 20, and map quality ≥ 30; these thresholds match those required for GATK and de novo variant calling. The following command was used to calculate the callable base pairs in each trio:

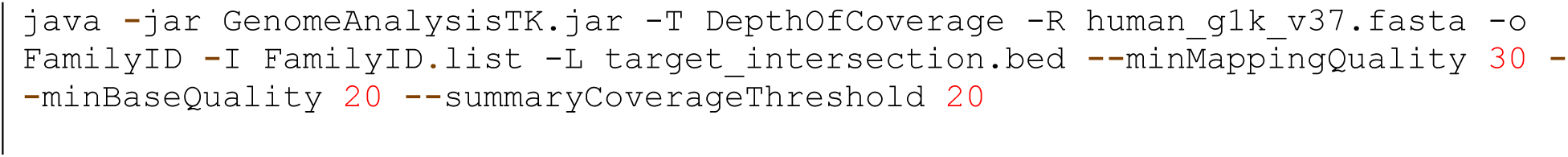

`FamilyID.list` contains names and locations of the three trio bam files. The .bed file contains the genomic intervals over which to calculate the callable base pairs. To calculate the coding callable base pairs (used for coding mutation rates, e.g. synonymous, nonsynonymous, missense, etc., see Table 1, Table S2), we used a bed file with intervals spanning the intersection of both capture array target intervals and the RefSeq coding intervals (32,027,823 bp total). To calculate all callable base pairs (used for the total coding + noncoding mutation rate, see “All” in Table 1), we used a bed file with intervals spanning the intersection of both capture array target intervals (33,973,867 bp total). The number of coding and total callable base pairs for every family passing is listed in Table S1.

### Contribution of de novo mutations to OCD risk

For every proband and control subject passing QC (obtained from Table S1), we made a file containing the number of Mis-D, LGD, and damaging (Mis-D + LGD) variants (obtained from Table S2), then calculated the haploid mutation rate per subject for each variant type. Haploid mutation rates were calculated by dividing the number of mutations by the twice the number of callable coding bases (“CallableExomeCoding” in Table S1).

The following R code was then used to calculate the percentage of cases with a mutation mediating risk and the percentage of mutations carrying risk, along with 95% confidence intervals for each. The number of de novo mutations per individual was calculated by multiplying the mutation rate by the size of the RefSeq hg19 coding exome (33,828,798 bp).

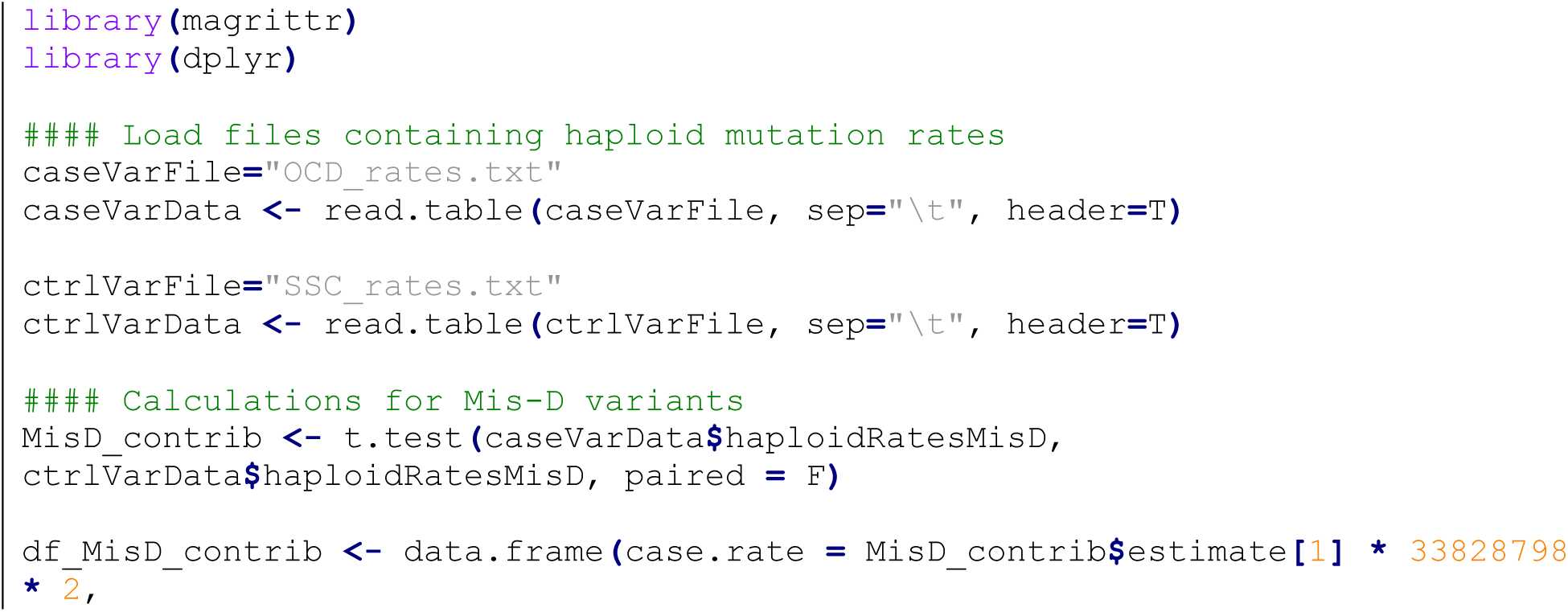

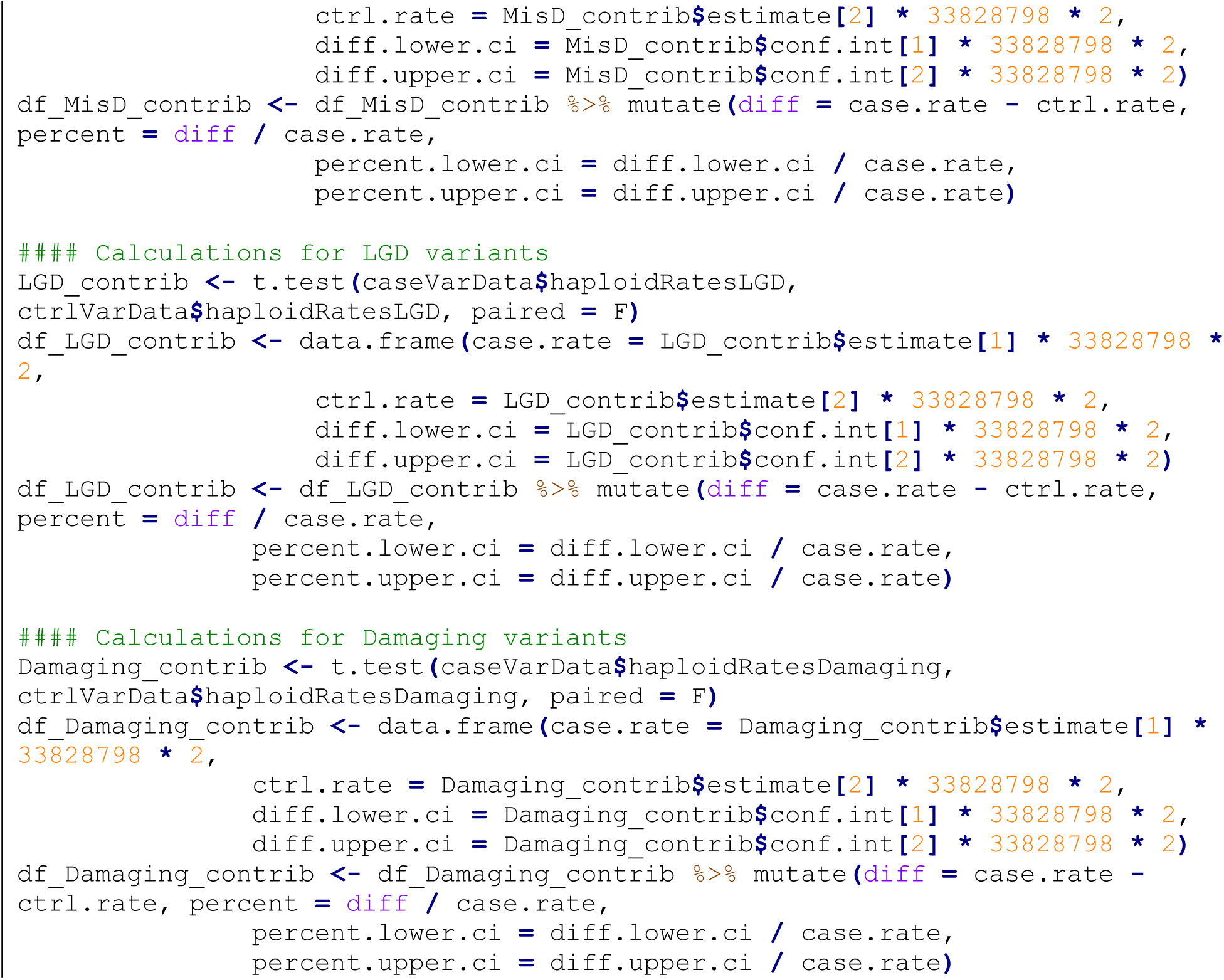

### Mutation rates for variant simulations

To perform subsequent maximum likelihood estimation and TADA analyses, we used published per gene de novo mutation rates from unaffected parent-child trios (1). For the control samples in our dataset, we calculated the proportion of the overall coding mutation rate that comprised LGD and Mis-D mutations, and then used these proportions to calculate the expected LGD and Mis-D mutation rate per gene.

The following R code was used to generate the mutation rate tables:

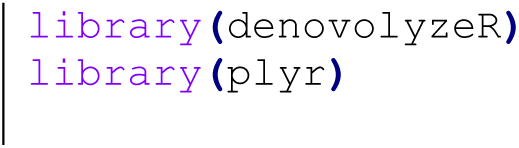

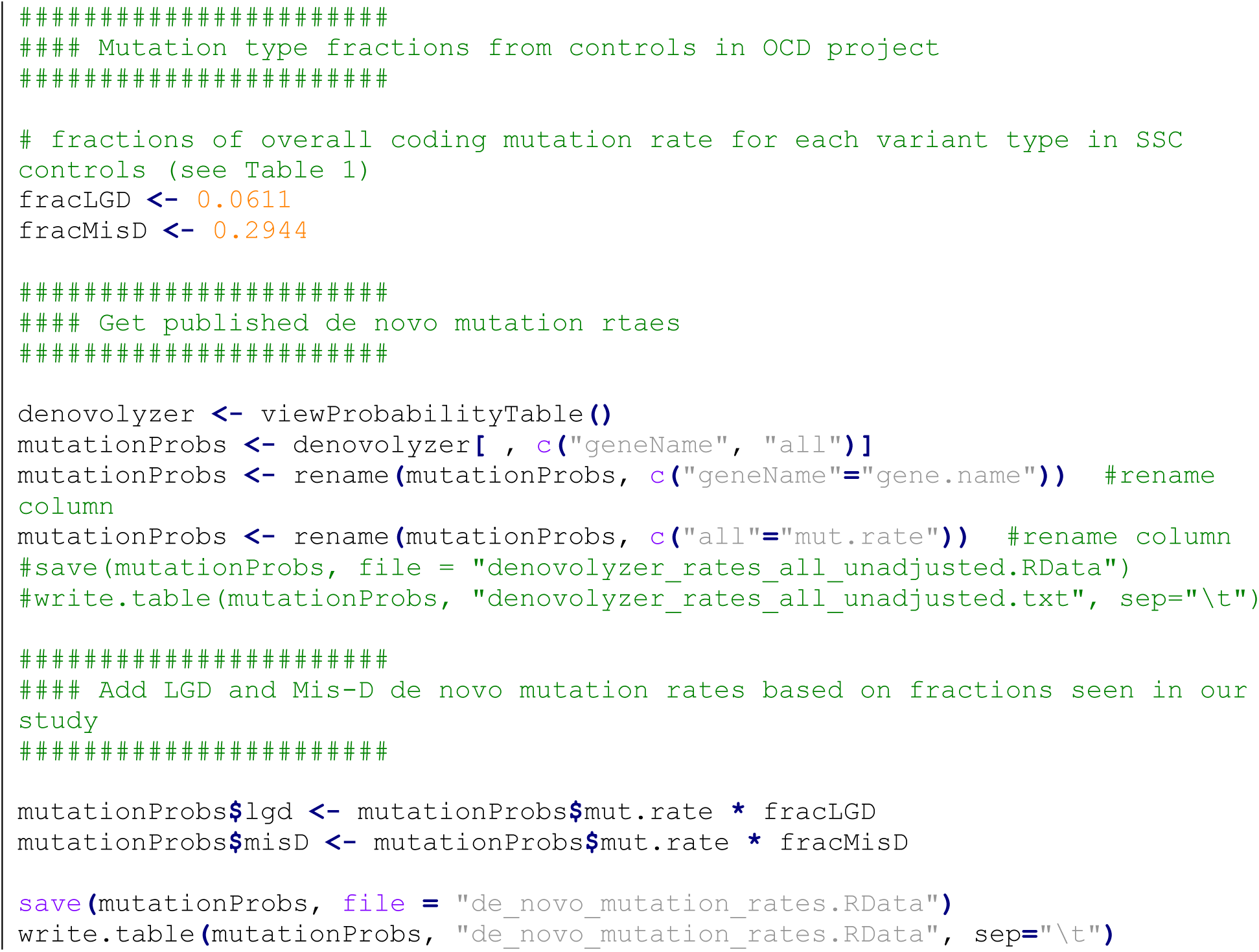

### TADA analysis

The enrichment of de novo LGD and Mis-D mutations in OCD raises the possibility that these classes of mutations target a set of genes that mediates OCD risk. To test this hypothesis, we used the transmitted and de novo association (TADA) test, a Bayesian model that can effectively combine data from de novo mutations, inherited variants in families, and standing variants in the population (via case-control cohorts) to significantly increase the power of gene discovery. In this study, we elected not to include the rare inherited exome variants because their confirmation rate is not known and their contribution to the TADA score is minimal given their lower relative risks (2, 3). Instead, we used a specialized version of TADA that analyzes only the de novo mutations from exome sequencing data, called TADA-Denovo (2). The code and documentation for this tool can be found here (TADA.v1.1.R; http://wpicr.wpic.pitt.edu/WPICCompGen/TADA/TADA_homepage.htm).

The TADA-Denovo test considers two types of variants, de novo LGD and de novo severe missense (those predicted by PolyPhen2-HDIV to be “probably damaging” to protein function, abbreviated as “Mis-D”). The main input is the number of de novo LGD and number of de novo Mis-D variants per gene. Additionally, mutation rates (mu) for all human genes are provided, based on Sanders et al. (4). The test analyzes each of these event types separately, then combines the evidence in a Bayesian fashion, weighing each type of mutation differently.

To compute the Bayes factors and p-values, TADA-Denovo requires the following parameters:

- n.family: the number of OCD parent-child trios passing QC (184)
- The fold-enrichment (λ) for each mutational class is calculated as follows, where X_case_ and X_ctrl_ are the number of mutations in cases and controls, respectively; S_case_ and S_ctrl_ are the number of synonymous mutations in case and controls, respectively: λ = (X_case_ / (X_ctrl_ · (S_case_/S_ctrl_))). As shown in the code below, we used the estimated coding variants per individual (Table 1) and multiplied by the number of cases or controls. Fold-enrichment for LGD was calculated as 2.09, and fold-enrichment for Mis-D was 1.46.
- The fraction of causal genes (π) is the estimated number of OCD risk genes (335, calculated by MLE as detailed below) divided by the number of RefSeq genes with mutational rates included in the TADA-Denovo algorithm. π = 335 / 19618 = 0.0171
- gamma.mean.dn: The average relative risk (γ) is related to the fold-enrichment (λ) and the fraction of causal genes (π) by the following equation: π · (γ-1) = λ-1. Solving for γ gives LGD γ = 64.74, Mis-D γ = 27.91).

Using these parameters, TADA-Denovo calculates the Bayes factors of all input genes. Next, it computes the p-value for each gene by generating random mutational data, based on each gene’s specified mutation rate, to obtain a null distribution of Bayes factors. We used 1,000 samplings of de novo mutations in each gene to determine null distributions. Finally, TADA-Denovo calculates an FDR q-value for each gene using a Bayesian “direct posterior approach.” A low q value represents strong evidence for OCD association. Genes with FDR < 0.3 are considered probable risk genes, and those with FDR < 0.1 are high-confidence risk genes.

The following R code performed the TADA-Denovo analysis:

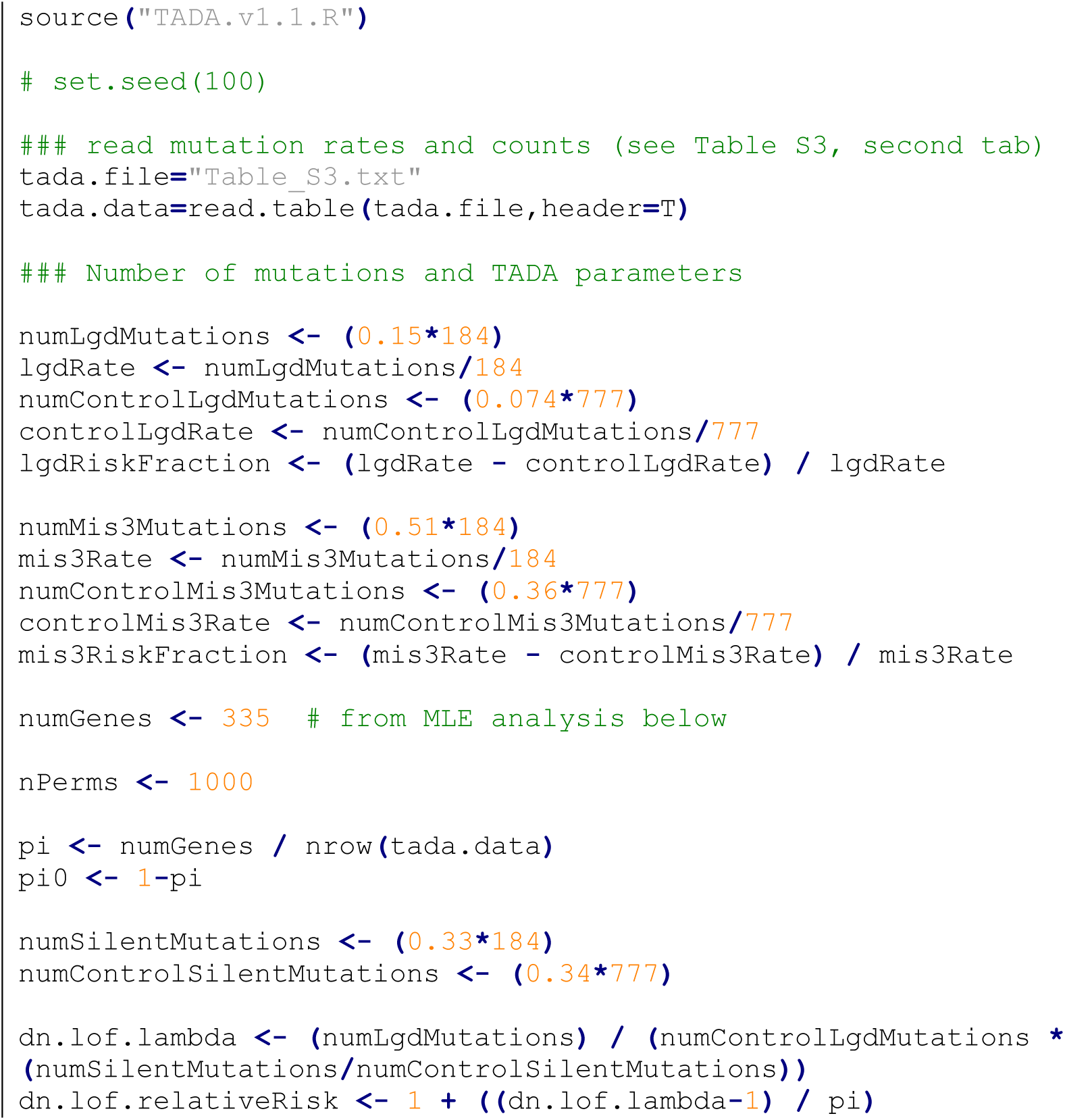

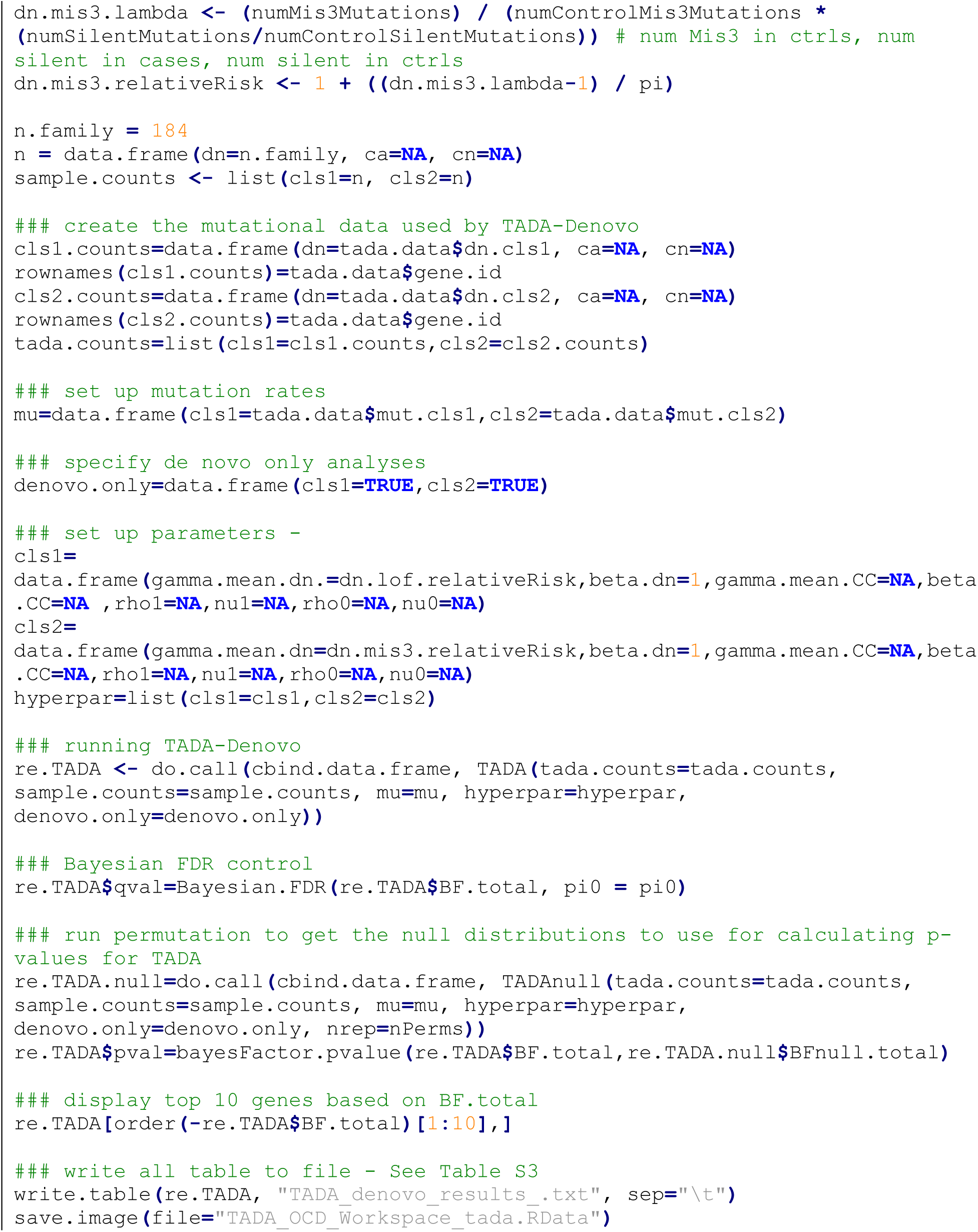

Applying this model to 184 OCD parent-child trios passing QC identifies two high confidence OCD risk genes (q<0.1) (Table 2, Table S3).

### Maximum Likelihood Estimation (MLE) method for estimating the number of OCD risk genes

The following R code was used to perform these calculations:

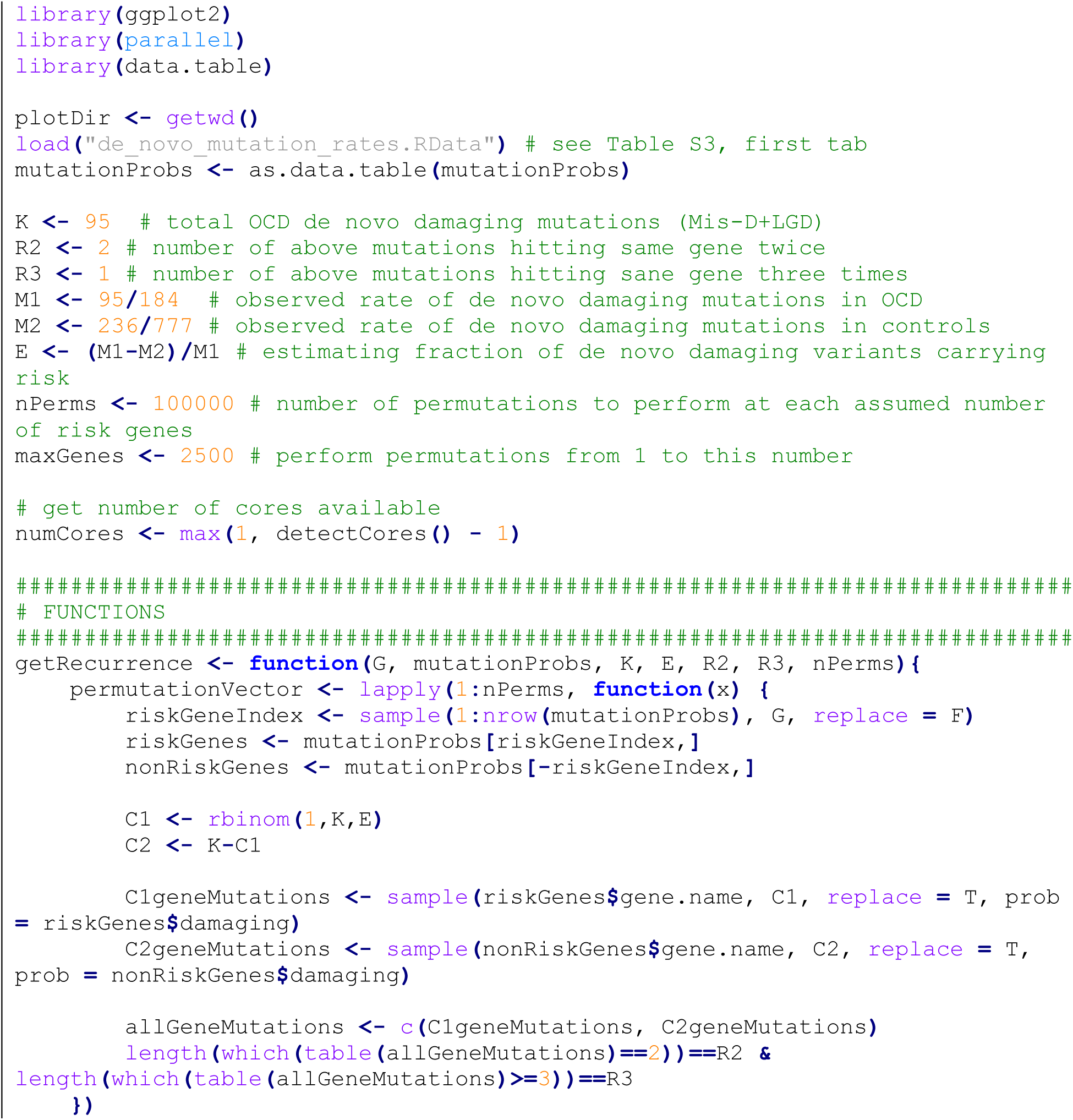

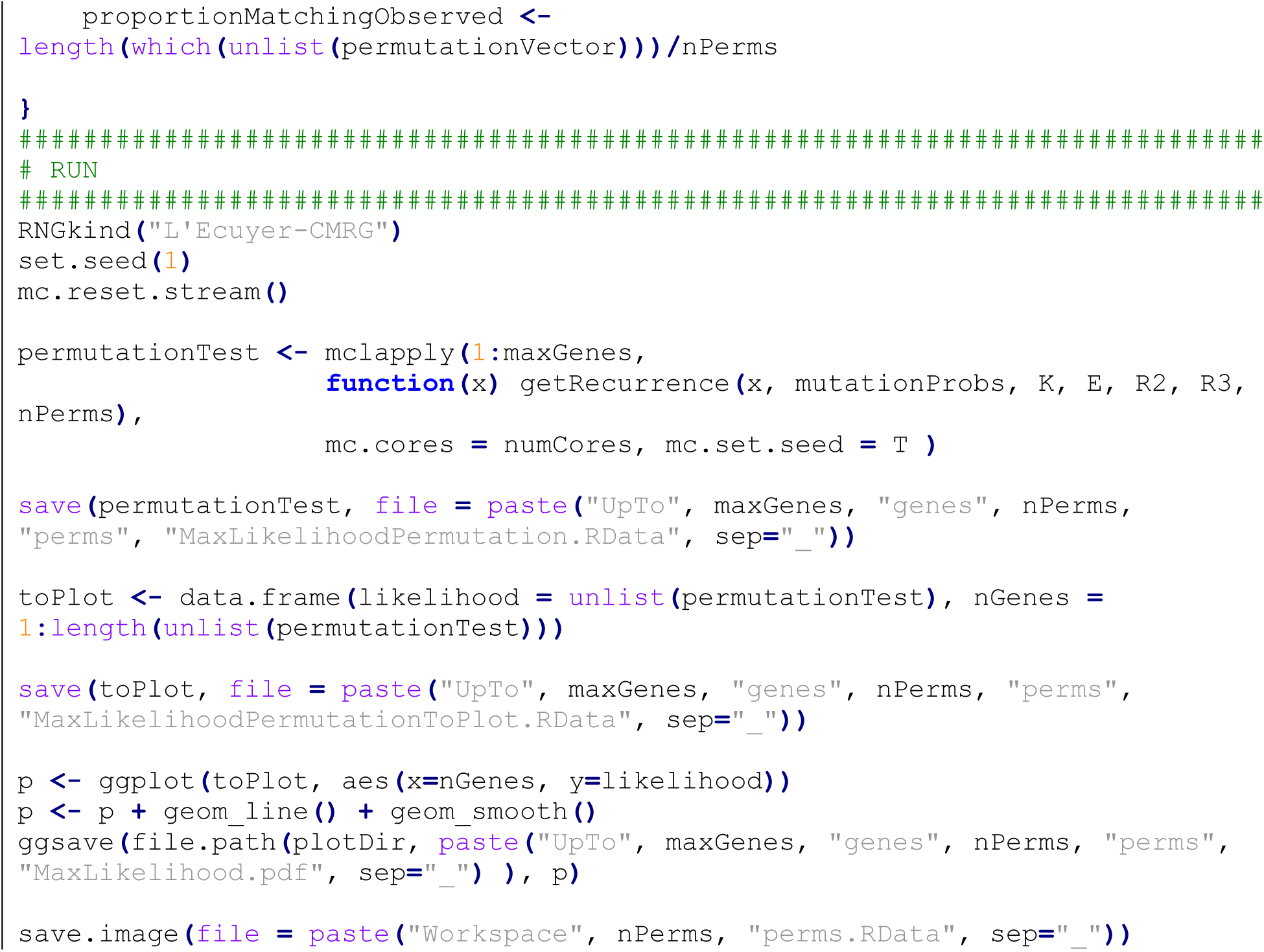

### “Unseen species” method for estimating the number of OCD risk genes

Following a method used previously in estimating the number of risk genes for autism spectrum disorder (4), we used the following R code to obtain a second estimate of the number of risk genes (C) in OCD and 95% confidence intervals.

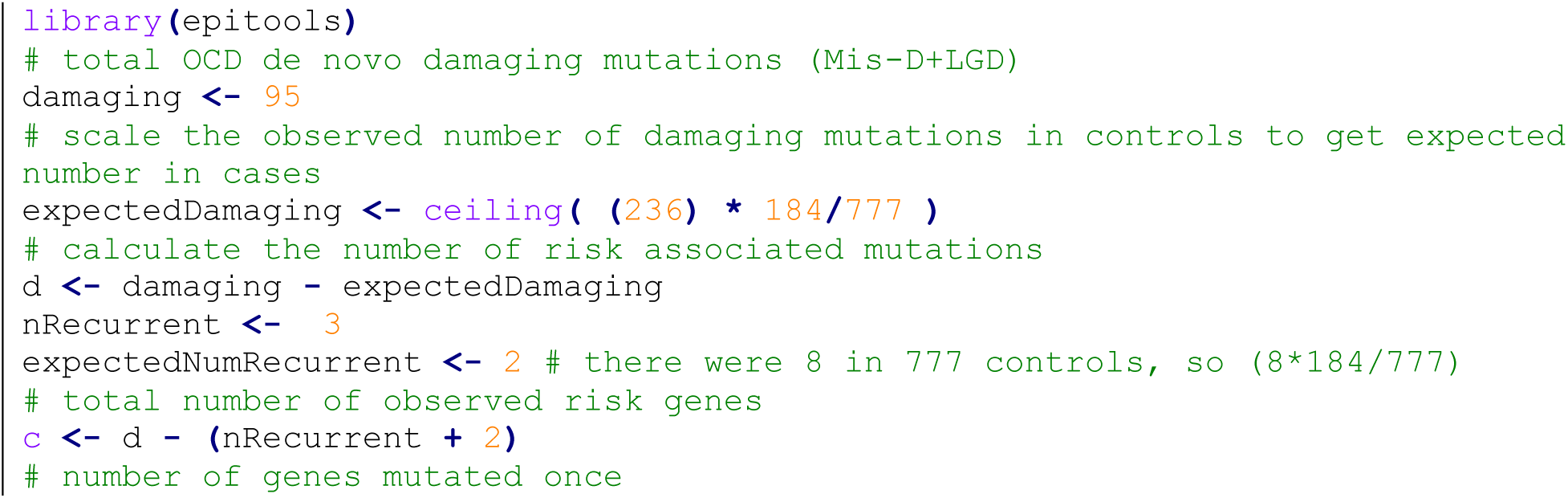

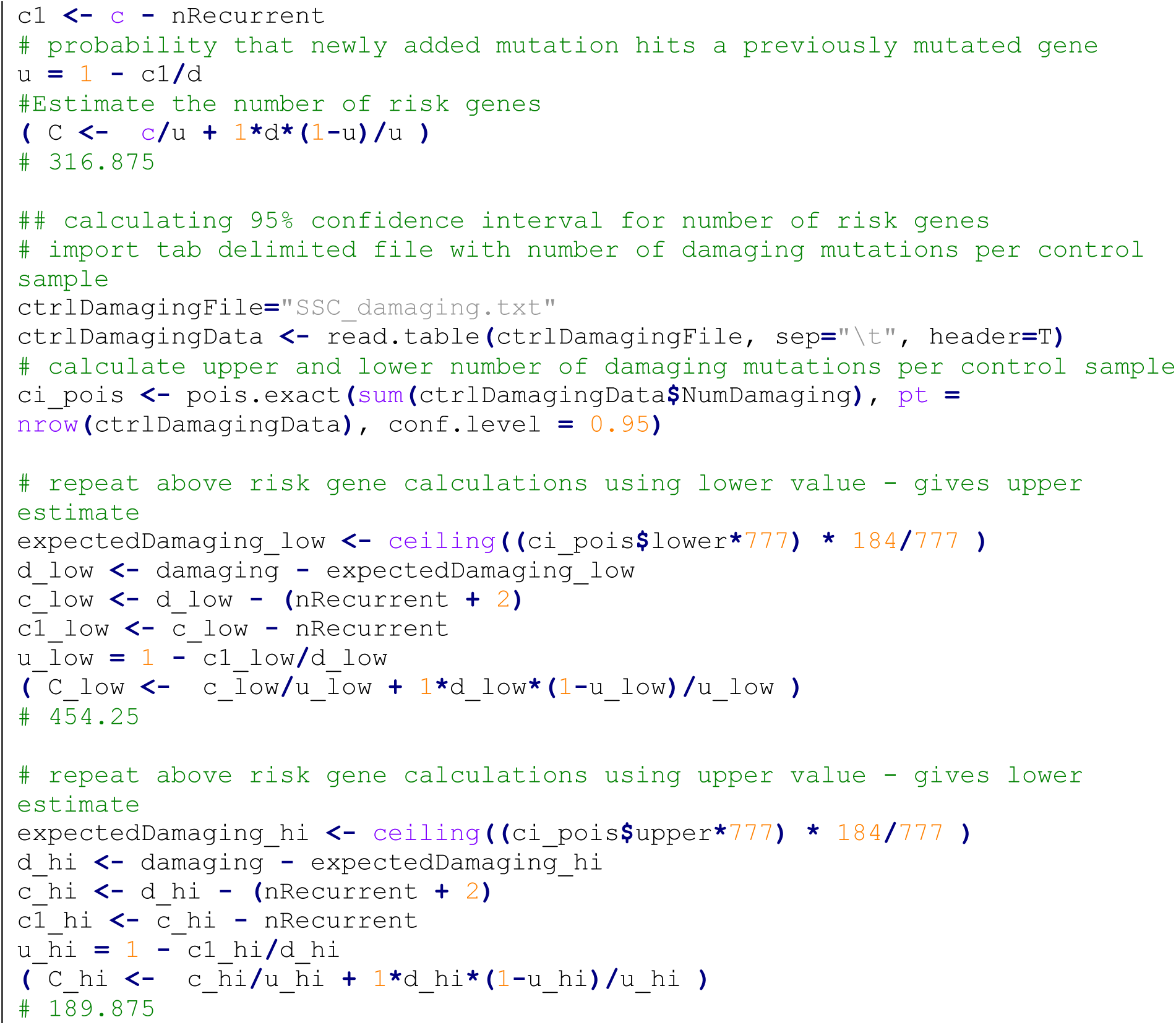

### Predicting the number of risk genes identified by cohort size

The following R code was used to perform these calculations:

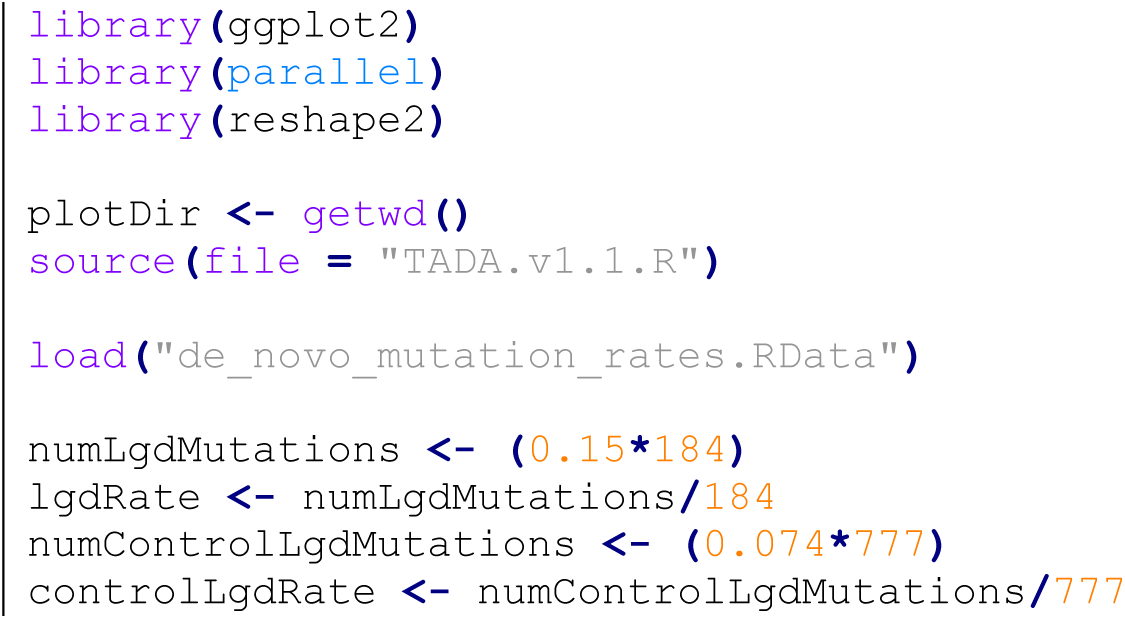

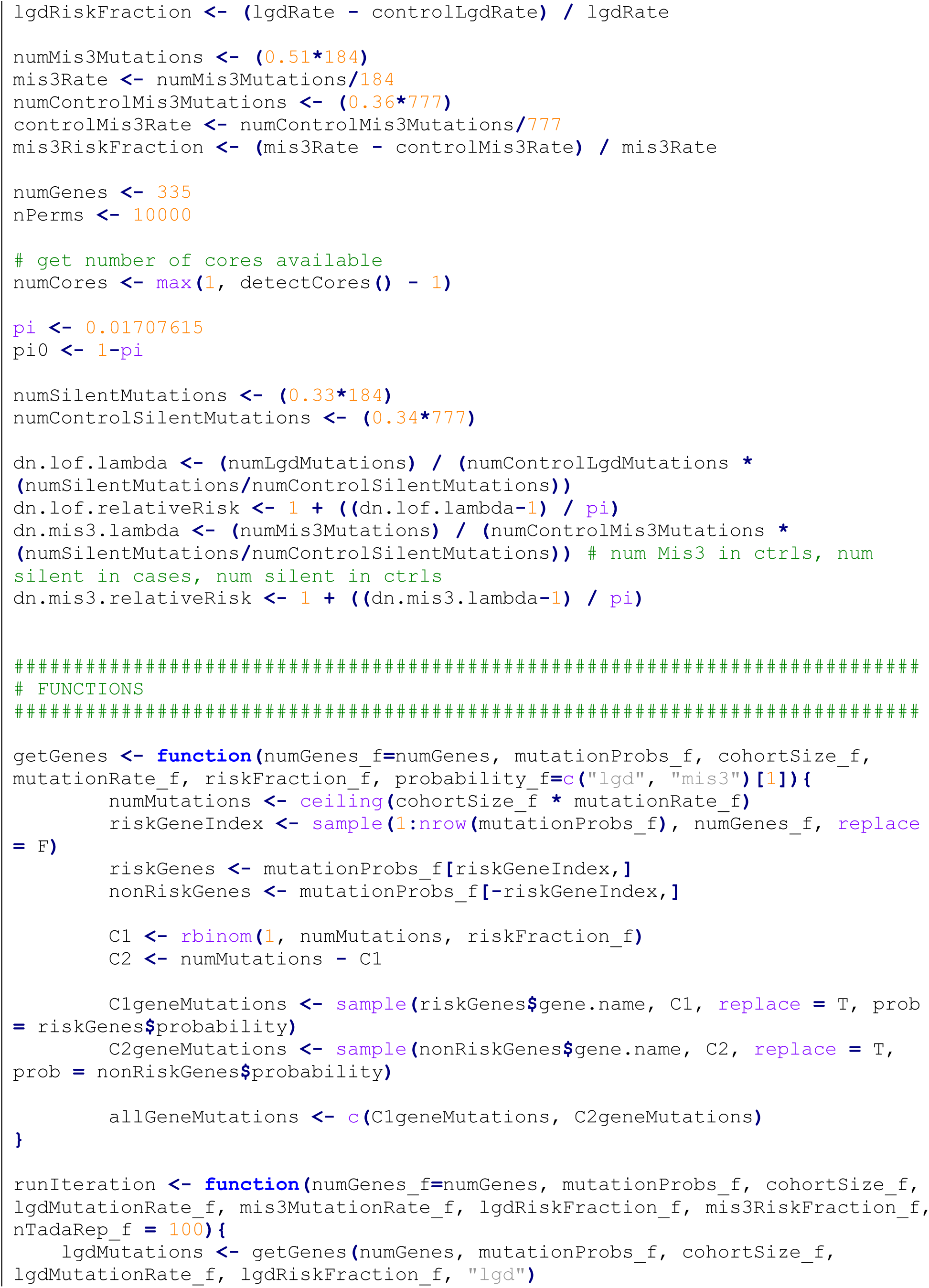

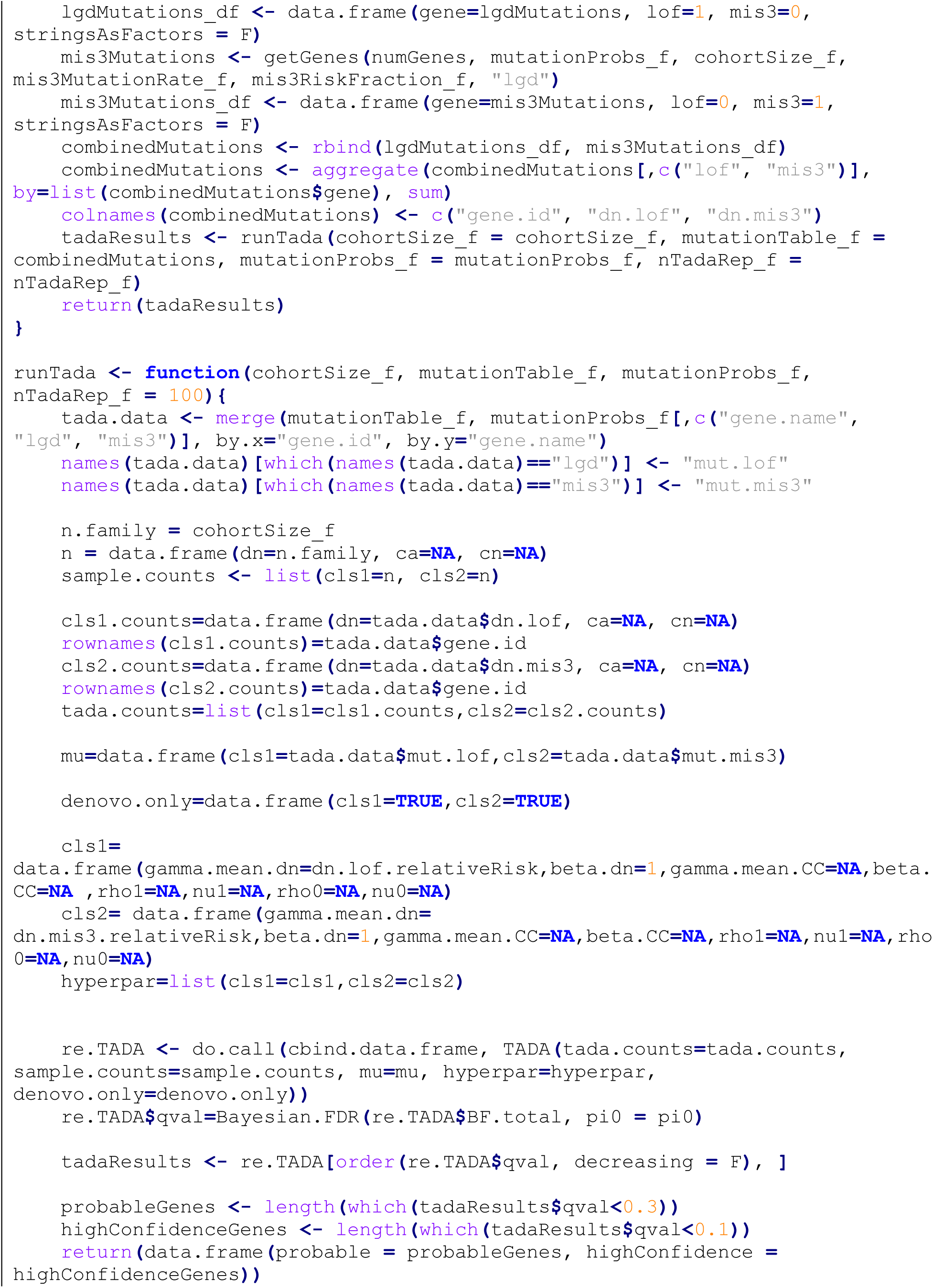

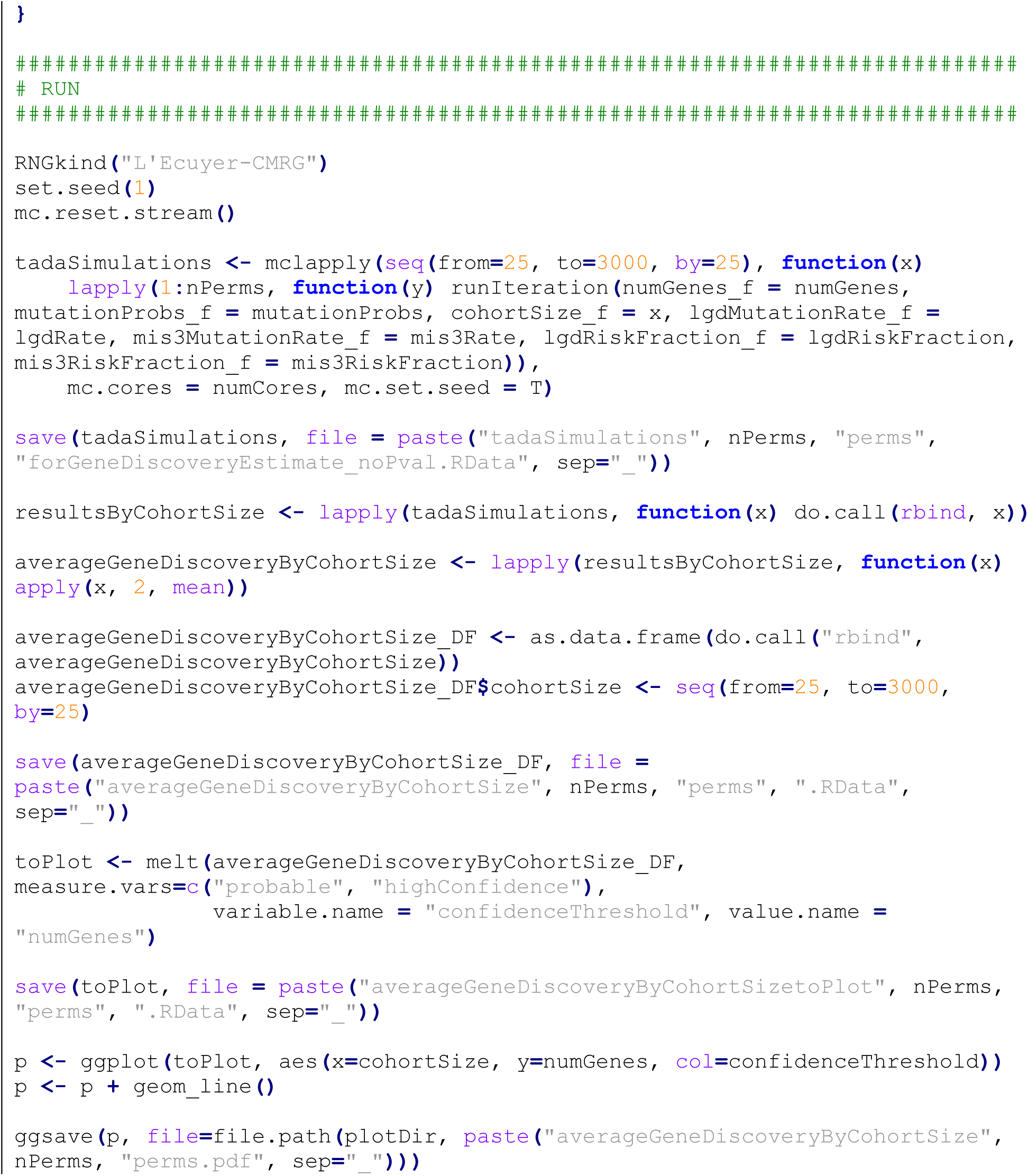

## Gene set overlap using DNENRICH

The following Linux commands were used to run the DNENRICH analysis:

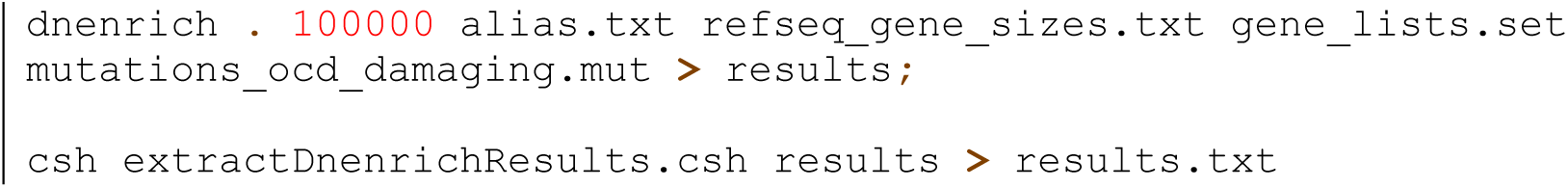

---

**Figure S1.**
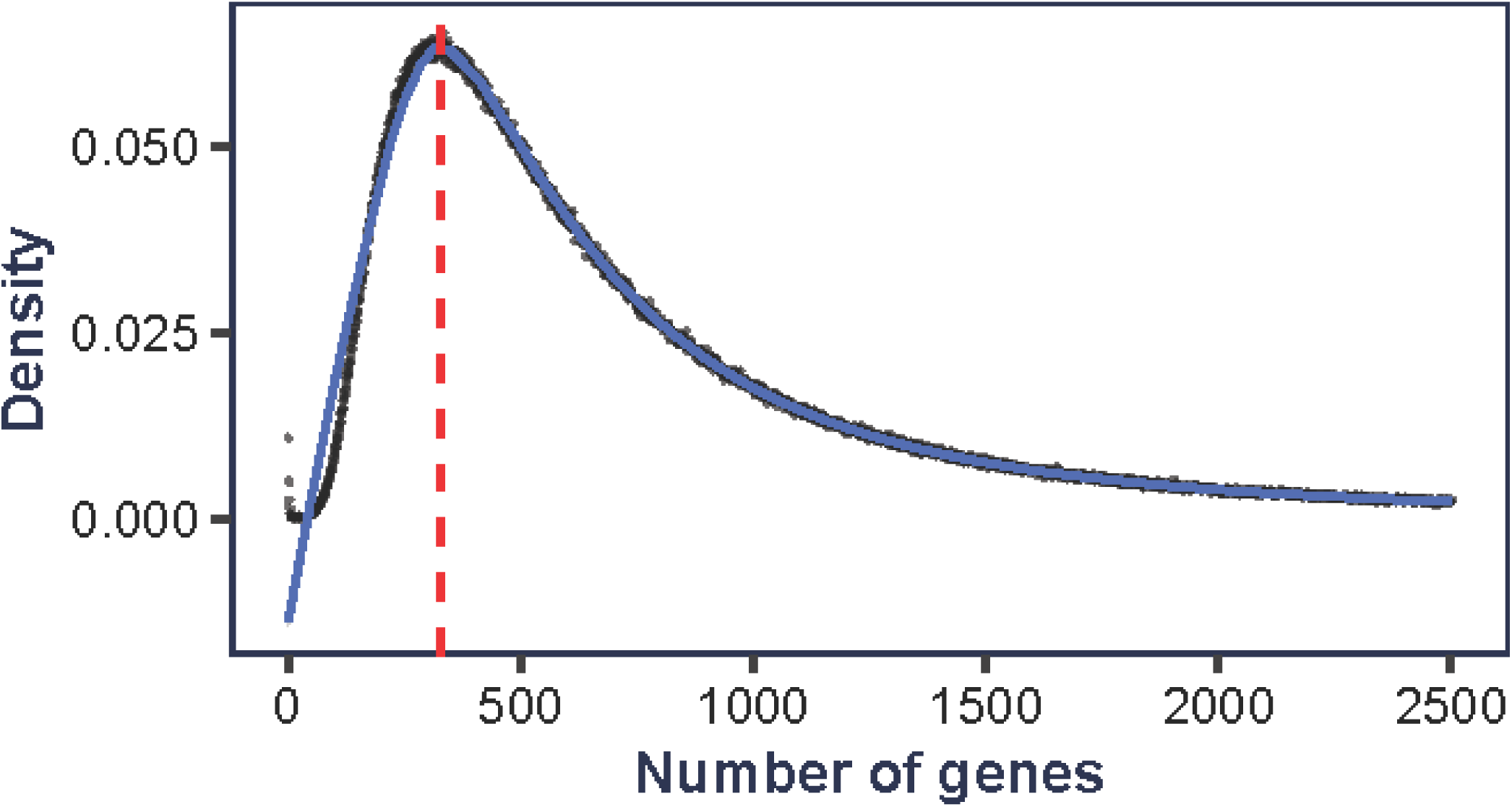
Maximum Likelihood Estimate (MLE) of number of OCD risk genes. Assuming each number of possible risk genes between 1-2,500, 100,000 simulations were conducted to determine the number of risk genes that yielded the closest agreement between our observed and simulated data. In each simulation, we generated 95 variants (the number of de novo damaging variants observed in our OCD sample), then randomly assigned a percentage of variants (determined by the fraction of de novo damaging variants estimated to carry OCD risk) to the risk genes, recording the frequency at which the number of genes with two and three recurrent variants matched the number observed in our study (2 and 1, respectively). This MLE method yields an estimate of 335 OCD risk genes (red vertical line), a number that is in close agreement with that from an alternative “unseen species” method (317 genes, 95% CI: 190-454).

**Figure S2.**
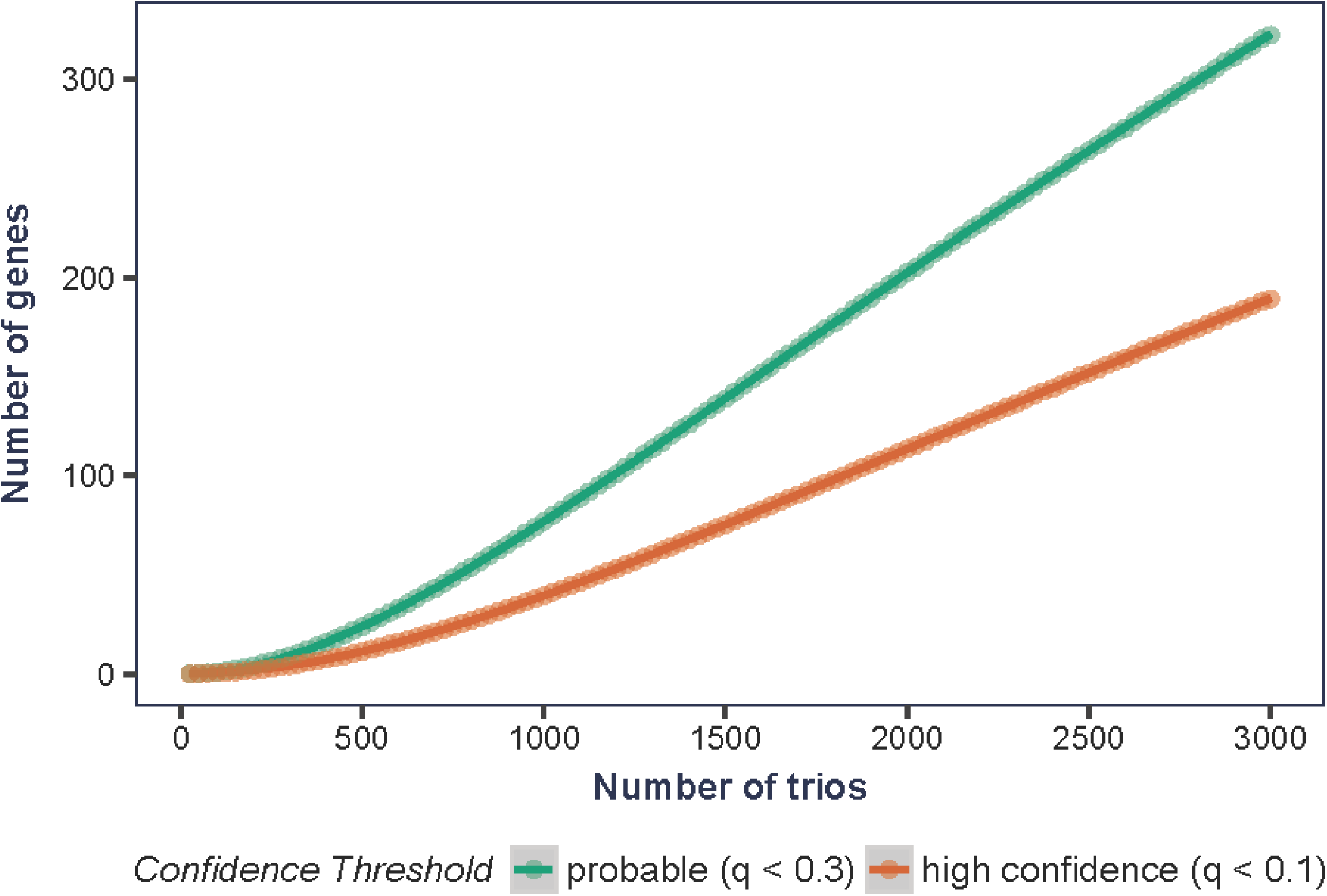
Gene discovery by number of trios sequenced. Using the MLE estimate of 335 risk genes, we estimated the number of probable (q<0.3) and high-confidence (q<0.1) risk genes that will be discovered as more OCD trios are sequenced. We performed 10,000 simulations at each cohort size from 25-3,000 trios, randomly generating variants and assigning to risk genes in agreement with the proportions seen in our data, then applying the TADA-Denovo algorithm.

**Figure S3.**
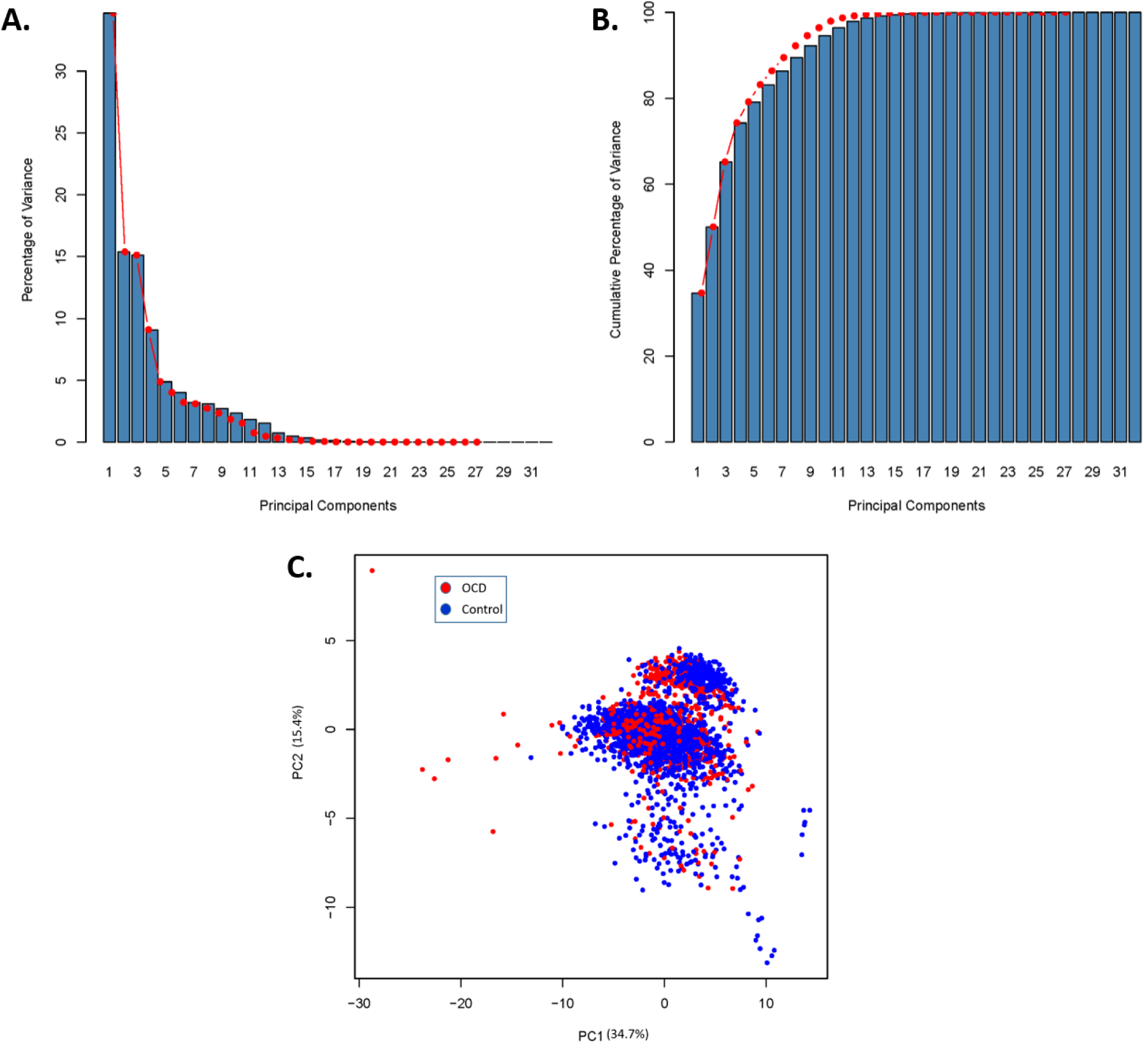
PCA scree and individual plots. Scree plots following Principal Components Analysis (PCA), showing (A) the percentage of variance captured by each of the first 32 principal components, and (B) the cumulative percentage of variance captured by these same components in the exome metrics data from cases and controls. The “elbow” of the scree plot is visualized to be around the 5^th^ principal component. This was confirmed by the Factominer R code function “estim_ncp()”. The first 5 PCs capture almost 80% of the variance, and this number of PCs was used to determine PCA outliers during quality control (see Table S1 and Supplementary Methods). (C) Individual plots for the first two principal components, based on PCA of exome sequencing quality metrics. OCD cases are plotted in red, and controls in blue. The first two PCs together capture 50.1% of the variance. R code to generate this data and figure are in Supplementary Methods, and individual PC factor values are in Table S1. This figure includes PCA outliers (>3 standard deviations from the mean in PCs 1-5), which were removed during quality control, prior to further analysis of case-control data.

**Figure S4.**
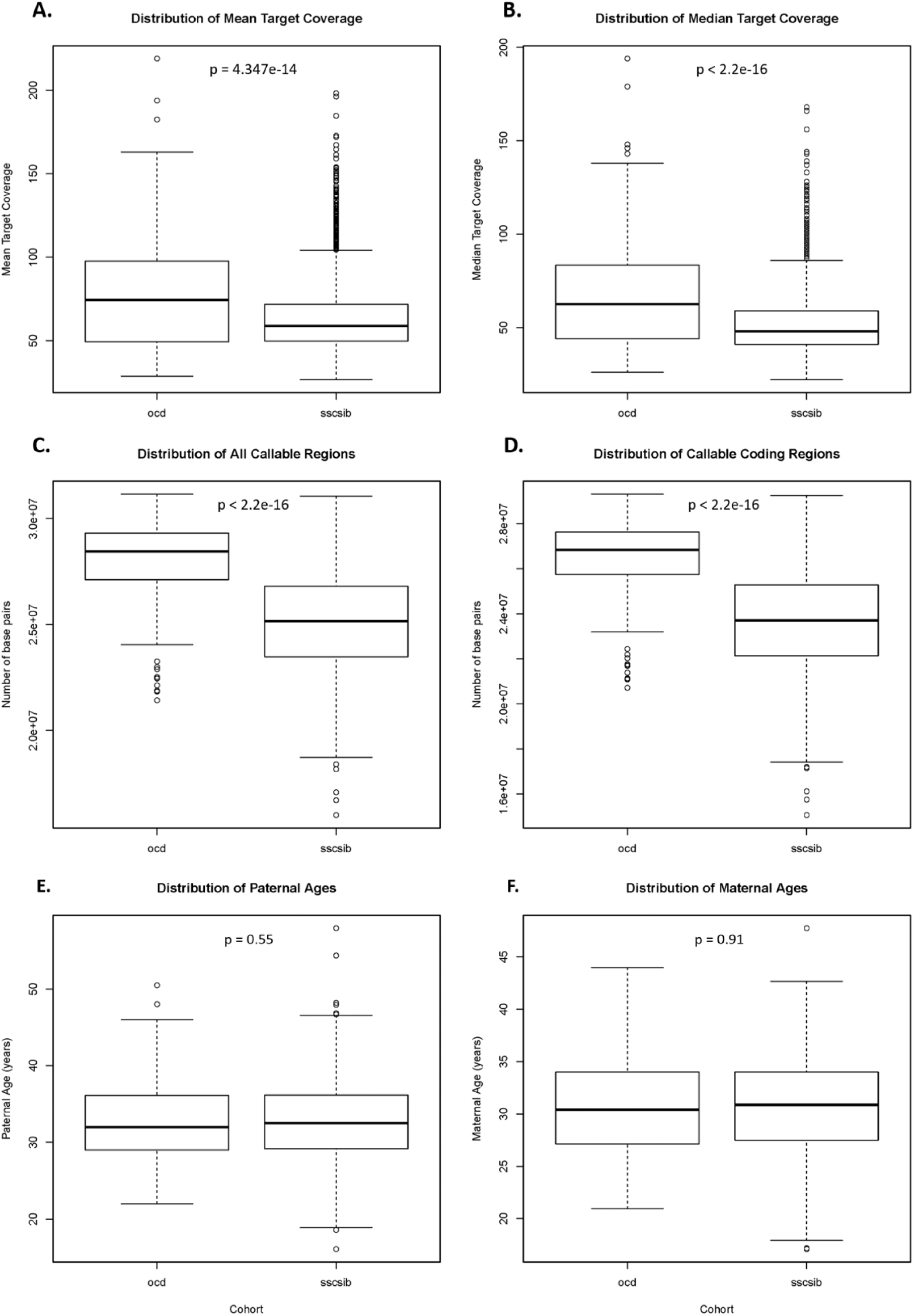
Sequencing coverage and parental age distributions. Distribution boxplots of values for (A) mean target coverage, (B) median target coverage, (C) number of base pairs in all “callable” regions, (D) number of base pairs in coding “callable” regions, (E) paternal age, and (F) maternal age for both OCD and control cohorts. For each cohort, the box extends from the first through third quartiles, and the horizontal line is at the second quartile (median) of the data. Whiskers extend to the largest non-outliers, and outlier data points are plotted individually. For each comparison, a p-value was calculated using a two-sided Wilcoxon rank sum test with continuity correction. Panels A-D show increased opportunity for variant calling in the OCD cohort, necessitating the use of de novo mutation rate comparisons within the callable exome, as explained in the main text and methods. Panels E-F show no significant difference in parental ages between case and control cohorts. Also see Table S1.

**Figure S5.**
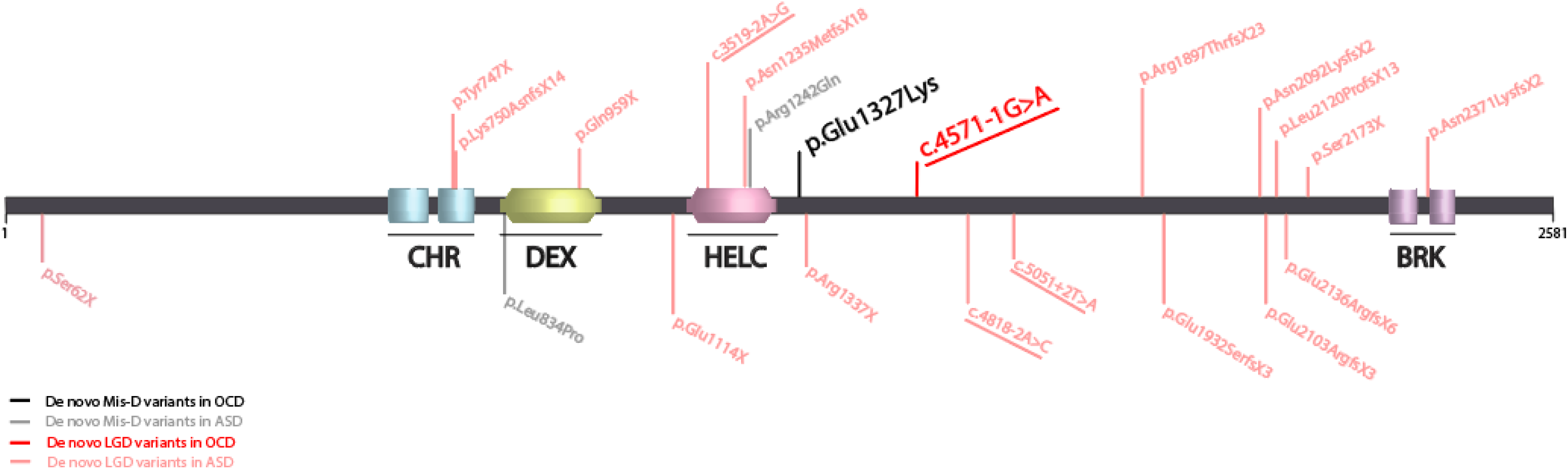
CHD8 variants in OCD and ASD. Two de novo likely gene disrupting (LGD, red) and damaging missense (Mis-D, black) variants identified in CHD8 among OCD probands are indicated. ASD-associated de novo LGD and Mis-D mutations reported in the Simons Foundation Autism Research Initiative (SFARI) database (accessed April 12, 2017) are also shown in muted colors. Only variants with identifiable allele or residue sequence positions in the SFARI database were included in the above protein diagram, and splice site variants across cohorts are indicated by the respective allele change and underlined. Annotated protein domains predicted with confidence by the Simple Modular Architecture Research Tool (SMART) are shown as follows: CHR, chromatin organization modifier domain (blue), DEX, DEAD-like helicases superfamily (yellow), HELC, helicase superfamily c-terminal domain (pink), and BRK, domain of unknown function associated with CHROMO domain helicases (purple).

**Figure S6.**
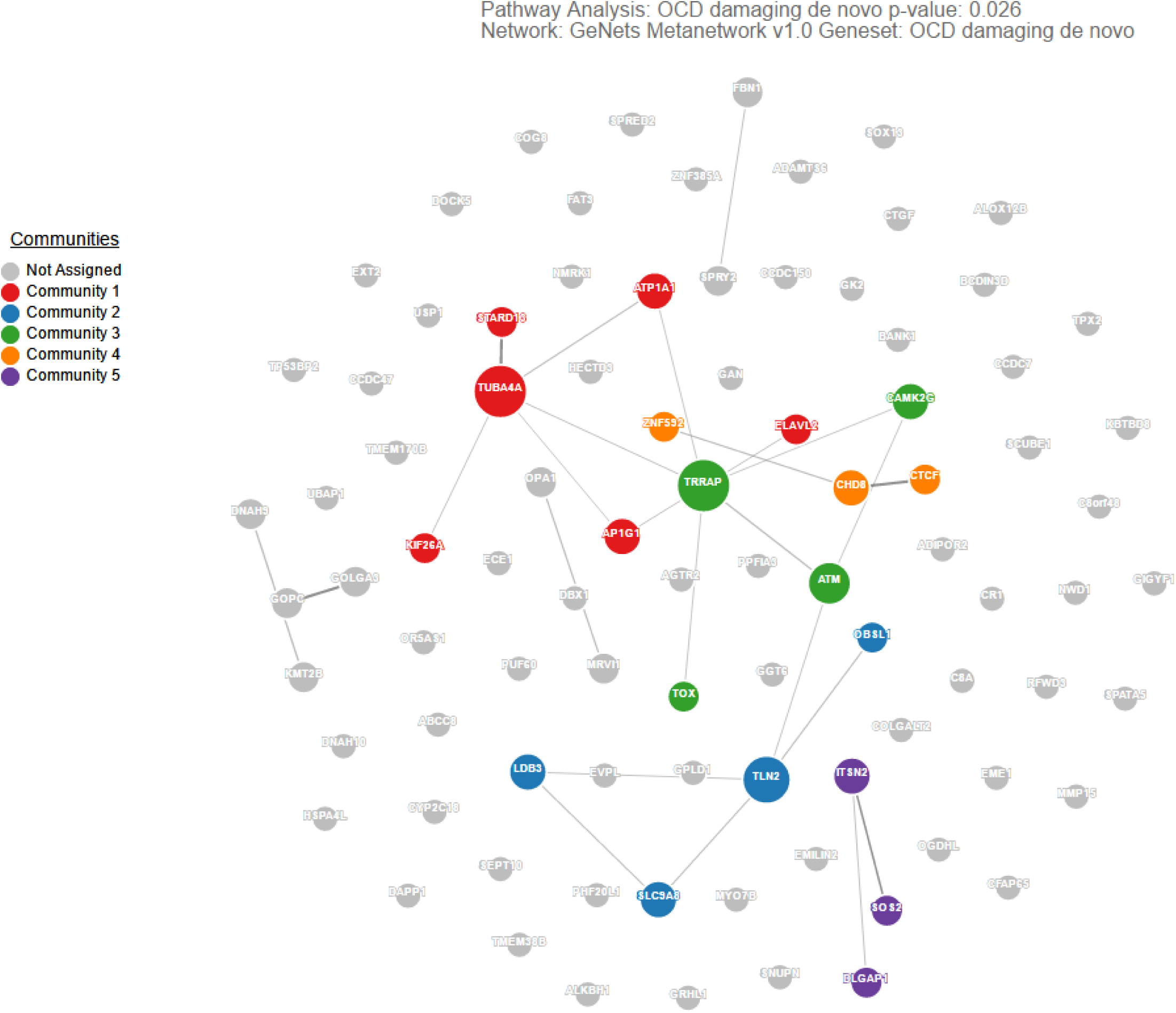
GeNets network analysis without candidates. Using the GeNets algorithm (https://apps.broadinstitute.org/genets), we mapped all 89 genes harboring de novo damaging mutations in OCD (excluding two genes, *TTN* and *CACNA1E*, found to harbor de novo damaging variants in control subjects) onto the GeNets Metanetwork v1.0 to determine whether they are functionally connected. The density of the mapped network (density = number of edges / number of possible edges) was greater than 95% of randomly sampled gene sets, indicating that the network is significantly more connected than random (p=0.026). In the network, node (gene) size is proportional to the number of connections. The color is assigned by community, defined as a gene set that is more connected to one another than to another group of genes. Results from this analysis are available in interactive form here: https://www.broadinstitute.org/genets#/visualize/58d9425ea4e00291af652379

## Supplemental Tables

**Table S1 - Phenotype, exome sequencing metrics, and principal components analysis.**

*(see “TableS1.xlsx”)*

First tab contains individual-level sample information (columns A-K), including family ID, individual ID, phenotype, cohort, collection site, gender, capture platform, size of “callable exome”, and parental age (years) at birth, where available. Column L lists reasons for any sample exclusions by quality control methods; “0” indicates that the sample was not excluded, and was included in subsequent analyses. Columns M-AH list individual sample sequencing metrics generated using PicardTools, and GATK DepthOfCoverage tools. Columns AI-AS list individual sample sequencing metrics generated using PLINK/SEQ (i-stats; https://psychgen.u.hpc.mssm.edu/plinkseq/stats.shtml). Columns B, M-AS were included in Principal Components Analysis (PCA). Third tab contains cohort-level metrics calculated using samples passing quality control. ±95% confidence intervals are given, when applicable. Fourth tab contains coordinates generated for each sample for the top 10 principal components following PCA. The code used to generate this data is included in Supplementary Methods. Using these coordinates, we removed trios with family members falling more than three standard deviations from the mean in any of the first five principal components; this information is contained in the fifth tab.

**Table S2 - Annotated de novo variants in OCD and controls.**

*(see “TableS2.xlsx”)*

Detailed information on all high confidence de novo variants in cases and controls. These variants were annotated using Annovar, based on RefSeq hg19 gene definitions. Column descriptions are provided in a separate tab of this file. A third tab provides the number of each de novo variant type per sample.

**Table S3 - Gene-level de novo mutation rates, variant counts, and TADA-Denovo results.**

*(see “TableS3.xlsx”)*

First tab contains de novo mutation rates used to perform subsequent maximum likelihood estimation (MLE) and TADA-Denovo analyses. The following mutation rates are listed for each gene: overall, likely gene disrupting (lgd), predicted damaging missense (misD), and all damaging (lgd + misD). These mutation rates were previously published (Ware et al., 2015) from unaffected parent-child trios. The code used to generate the mutation rate table is provided in Supplementary Methods. Second tab contains the input file for the TADA-Denovo algorithm. Gene-level expected mutation rates for LGD (“mut.cls1” column) and Mis-D variants (“mut.cls2” column) are listed, along with their respective observed mutation counts in our OCD data (“dn.cls1” and “dn.cls2”, respectively). Code for running TADA-Denovo is given in Supplementary Methods. Third tab contains the final output results from TADA-Denovo code provided in Supplementary Methods. Genes harboring more than one damaging de novo (LGD or Mis-D) variant in OCD probands are highlighted in yellow (*SCUBE1, CHD8, TTN*). Two of these genes (*SCUBE1* and *CHD8*) exceeded thresholds for being considered a probable (qval < 0.3) or high confidence (qval < 0.1) risk gene.

**Table S4 - DNENRICH gene lists and results.**

*(see “TableS4.xlsx”)*

See Supplementary Methods for details of DNENRICH analysis. First tab contains input for DNENRICH analysis. Each row represents a de novo damaging mutation in an OCD proband. Second tab contains the input gene lists to determine enrichment for our OCD damaging de novo mutations. Third tab contains final results output from DNENRICH. Significantly enriched gene sets are highlighted.

**Table S5 - GeNets network connectivity analysis results.**

*(see “TableS5.xlsx”)*

Complete results from GeNets network analysis of de novo damaging variants found in OCD probands. First tab contains summary statistics of the resulting network, considered both with and without nearby predicted candidate genes. Second tab contains the input gene list and the candidate genes predicted by the network analysis. Third tab groups genes (without predicted candidates) into nearby “communities” that are more connected with each other than their neighbors. Fourth tab contains network edges without the predicted candidates. Fifth tab contains gene community groupings, including predicted candidates. Sixth tab contains network edges including predicted candidates. See Methods for further details of this analysis.

**TableS6 - MetaCore and Ingenuity Pathway Analysis (IPA) gene enrichment analysis results.**

*(see “TableS6.xlsx”)*

Complete results from Metacore (first tab) and IPA (second tab) gene enrichment analyses, with p-values calculated by each analysis algorithm. See Methods for details of these analyses.

